# *Mimosa* species endemic to acidic soils in central Brazil are nodulated by a high diversity of *Paraburkholderia* genotypes, but widespread species are nodulated by *Paraburkholderia*, *Cupriavidus* or *Rhizobium* depending on soil characteristics

**DOI:** 10.1101/2024.06.11.598400

**Authors:** Luc Rouws, Alexandre Barauna, Chrizelle Beukes, Janaina R. C. Rouws, Sergio Miana de Faria, Eduardo Gross, Fabio Bueno dos Reis Junior, Marcelo F. Simon, Marta Maluk, David W. Odee, Stephanie Fordeyn, Gregory Kenicer, J. Peter W. Young, Veronica M. Reis, Jerri Zilli, Euan K. James

## Abstract

Neotropical Beta-rhizobia have a particular affinity to the large legume (*Fabaceae*) genus *Mimosa* and some of its relatives in the tribe Mimosae of the Caesalpinioideae subfamily. However, little is still known about the ecology of this interaction, especially the relationship between the rhizobia of “widespread” pan-tropical *Mimosa* species like *M. pudica* and the rhizobia that nodulate endemic *Mimosa* species that are very restricted in their habitats. The objective of this study was to examine the microsymbionts of *Mimosa* spp. and some other mimosoids in climates ranging from tropical to subtropical, humid to semi-arid, with varied soil characteristics and altitudes, with the aim of testing the hypothesis that widespread species have more cosmopolitan symbiont preferences than endemic ones. Nodules were sampled from more than 40 *Mimosa* spp. and related taxa in eleven Brazilian states, many endemics or biome-restricted, but particular attention was paid to sample nodules from the widespread species *M. pudica* at all locations. The *Mimosa* symbionts comprised 19 potential 16S rRNA and *recA* groups at the species level, with 16 belonging to the genus *Paraburkholderia*, including six lineages that may represent new species. The remaining genotypes consisted of 14 strains in two lineages of *Cupriavidus* that were mainly isolated from *M. pudica* growing at low altitudes, plus a single lineage of *Rhizobium* also from *M. pudica*. It is concluded that a high diversity of *Paraburkholderia* strains dominate as symbionts of *Mimosa* in the acidic soils of its main center of radiation in Central Brazil but that *Cupriavidus* and *Rhizobium* comprise a significant minority of symbionts of widespread *Mimosa* spp., especially *M. pudica*, in lowland or disturbed areas with less acidic soils. *Mimosa* symbiont diversity is thus driven either by edapho-climatic characteristics for widespread species and/or by co-evolution of the symbiotic partners for endemic species.

## 1. Introduction

Rhizobia are a polyphyletic group of soil-borne bacteria that form symbioses with leguminous plants (Fabaceae) and the non-legume *Parasponia* (Cannabaceae) (Sprent, 2009; Reeve et al. 2015; Sprent et al. 2017; Ardley and Sprent 2021). They induce nodule formation on the roots (and occasionally the stems) of their hosts and supply them with nitrogen-rich compounds obtained through a process called Biological Nitrogen Fixation (BNF) in which atmospheric N_2_ (dinitrogen) is converted to NH_3_ (ammonia) within the nodules via the enzyme nitrogenase (Remigi et al., 2016; Sprent et al. 2017). The currently known rhizobia belong to the *Alphaproteobacteria* and *Betaproteobacteria* bacterial classes and are commonly referred to as alpha- and beta-rhizobia, respectively (Gyaneshwar et al. 2011; Peix et al. 2015; Sprent et al. 2017; Ardley and Sprent 2021). The Alpha-rhizobia consist of *Rhizobium* and *Bradyrhizobium* plus several other genera mainly in the order *Hyphomicrobiales* (formerly *Rhizobiales*). In contrast, the Beta-rhizobia consist of only three genera to date, all in the order *Burkholderiales*: *Paraburkholderia*, *Cupriavidus* and *Trinickia* (Chen et al. 2001, 2003a, b; Beukes et al. 2017; da Silva et al. 2012; Andrews and Andrews 2017; Estrada de los Santos et al. 2018; Ardley and Sprent 2021).

Among the six legume sub-families, only two of them, the Caesalpinioideae and the Papilionoideae, harbor nodulating genera. Nodulation is almost ubiquitous in the Papilionoideae (97% of genera nodulated), but much rarer in the Caesalpinioideae (33%), with the notable exception of the Mimosae tribe (93%) nested within it (Sprent, 2009; Sprent et al. 2017; de Faria et al. 2022). Alpha-rhizobia, such as *Rhizobium* and *Bradyrhizobium*, nodulate several genera across both nodulating sub-families, but nodulation by the Beta-rhizobia is so far reported in only a few legume genera, most particularly with some neotropical members of the Mimosae tribe, but also with a range of endemic papilionoid legumes in the Fynbos biome at the tip of South Africa (Elliott et al. 2007b; Garau et al. 2009; Gyaneshwar et al. 2011; Beukes et al. 2013; Howieson et al. 2013; Liu et al. 2014; Lemaire et al. 2015, 2016; Dludlu et al. 2018a, b; Mavima et al. 2021, 2022). Arguably the most widely reported legume genus with which all three known Beta-rhizobial genera can nodulate is the large mimosoid genus *Mimosa* (Chen et al. 2001, 2003a, b, 2005a, b; Barrett and Parker 2005, 2006, Elliott et al. 2007a, 2009; Bontemps et al. 2010; dos Reis Junior et al. 2010; Gyaneshwar et al. 2011, Mishra et al. 2012; Lammel et al. 2013, 2015; Platero et al. 2016; Estrada de los Santos et al. 2018; Paulitsch et al. 2019a, b, 2020a, b; Mavima et al. 2021; Klepa et al. 2021; Dias et al. 2021), but they are also reported to nodulate other mimosoid genera, such as members of the “Piptadenia Group” (Taulé et al. 2012; Bournaud et al. 2013, 2017), and the large neotropical genus *Calliandra* (Silva et al. 2018; Zilli et al. 2021).

Current data suggest that except for the South African Fynbos biome (see earlier), Beta-rhizobia originated in Central and South America and co-evolved with the ancestors of mimosoid genera over the last 50 million years (Bontemps et al. 2010; Gyaneshwar et al. 2011); much more recently (last 500 years), some Beta-rhizobia were globally dispersed by the human-mediated transport of soil and neotropical plants from the Americas to much of the tropical world, such as South East Asia and Australia, mostly with *Mimosa pudica*, *M. diplotricha*, and *M. pigra*, all of which are now established as highly invasive weeds in these environments (Chen et al. 2001, 2005b; Parker et al. 2007; Elliott et al. 2009; Gyaneshwar et al. 2011; Klonowska et al. 2012, 2018; Liu et al. 2012, 2020; Andrus et al. 2012; Gehlot et al. 2013; Melkonian et al. 2014). Over the last two decades, most studies have identified the Beta-rhizobial genus *Paraburkholderia* as the predominant nodule-forming bacteria of *Mimosa* spp., mainly occurring in acidic tropical soils in Central and South America, indicating soil pH as a determinant factor for the selection of *Paraburkholderia* as a symbiont (Chen et al. 2005a; Bontemps et al. 2010; dos Reis Junior et al. 2010; Mishra et al. 2012; Pires et al. 2018). It is supported by studies from other types of soils, *e.g.*, *Mimosa* spp. growing in basic soils in India and Mexico preferred alpha-rhizobial symbionts (Gehlot et al. 2013; Bontemps et al. 2016), as did those in unusually high pH soils in central Brazil (Pires et al. 2018), whereas those in high-pH soils in Uruguay nodulated exclusively with neutral-alkaline-preferring *Cupriavidus* spp. (Platero et al. 2016; de Meyer et al. 2015a).

*Mimosa* in the tribe Mimosae is one of the largest genera in the subfamily *Caesalpinioideae* with more than 600 species (LPWG, 2023); almost 60% of its species (app. 380) are native or endemic to Brazil (Simon and Proença 2000; Dutra et al. 2020), with a secondary center of radiation in Mexico (Simon et al. 2011). *Mimosa* species extend from northern Argentina in the south to the southern USA in the north, where they grow in diverse soils and climates, but are particularly endemic in higher altitude locations, *e.g.*, in the Cerrado and Caatinga biomes of Brazil (Simon and Proença 2000; Simon et al. 2009, 2011; dos Reis Junior et al. 2010). The ecology of the symbiosis between *Mimosa* and rhizobia is relatively understudied, especially considering the environmental factors affecting the symbiosis with various rhizobial types of such a widely distributed legume genus. However, earlier work by Bontemps et al. (2010) involving a large-scale survey of symbionts from 47 *Mimosa* species native or endemic to the Brazilian Cerrado, Caatinga and Pantanal biomes suggested the co-evolution of native Brazilian *Mimosa* species with nodulating symbionts in the betaproteobacterial genus *Burkholderia*. Bontemps et al. (2010) identified seven Species Complexes (SC) based on the 16S rRNA-*recA* sequences of 143 strains; these same deep-branched SC were congruent with the lineages recovered in the phylogenies of the symbiosis-essential genes, *nodC* and *nifH*, suggesting little horizontal gene transfer (HGT) had occurred. It was thus concluded that their partnership with *Mimosa* was “ancient” rather than inherited recently from alpha-rhizobia via HGT. Bontemps et al. (2010) also noted that one of the larger SC in their study, SC5, predominated as symbionts of mainly endemic species growing above 1000 m, *e.g.*, in the Chapada dos Veadeiros in the state of Goias (GO) and the Chapada Diamantina in the state of Bahia (BA), both locations wherein *Mimosa* shows high levels of diversification and endemism (Simon and Proença 2000; Simon et al. 2011), and hence further concluded that elevation also played a role (directly or indirectly via the host) in the selection of symbionts by *Mimosa* species endemic to highland regions.

Since it was published, almost all the symbionts in the study of Bontemp et al. (2010) have been largely moved to a new genus *Paraburkholderia* (Sawana et al. 2014), with a small number of strains being incorporated into another new genus, *Trinickia* (Estrada de los Santos et al. 2018). In terms of the current taxonomy of *Burkholderia sensu lato*, the *Burkholderia* SCs identified by Bontemps et al. (2010) contain the following species: SC1 (*Trinickia symbiotica*), SC2 (*P. sabiae*, *P. caribensis*, *P. phymatum*), SC3 (*P. diazotrophica, P. franconis*), SC4 (*P. mimosarum*), SC5 (*P. nodosa* and *P. guartelaensis*), SC6 (*P. tuberum* sv. mimosae, now moved to *P. atlantica* and *P. youngii*), and SC7 (*P. phenoliruptrix*). It should be noted, however, that all the SCs also contain several strains that are not yet allocated to a formally described species.

The overall objective of this study was to test the hypothesis that widespread *Mimosa* species have more cosmopolitan symbiont preferences than endemic ones in the main center of this legume genus, Brazil. Towards this aim, the occurrence of nodulating *Mimosa* spp. was surveyed across their native range in several states of Brazil, encompassing various biomes. Rhizobial diversity is almost certainly at least partly related to soil characteristics (Elliott et al. 2009; Bontemps et al. 2016; Pires et al. 2018; Soares-Neto et al. 2022), given the solid biogeographical relationships between *Mimosa* diversity and its native environments. However, it is difficult to establish which factors other than co-evolution are involved in the selection of symbionts by many *Mimosa* spp. because the plants are either highly endemic or biome-restricted (Simon and Proença 2000; Simon et al. 2011). Therefore, in order to mitigate the effects of co-evolution, the present study had a particular emphasis on the symbionts of widespread/invasive species that colonize many types of soils and which are “promiscuous” in terms of their rhizobial preferences, *e.g.*, *M. pudica* (see aforementioned references). The symbionts from these were compared to type strains, but also to reference strains from previous studies of the symbionts of *Mimosa* (and other mimosoids, such as *Calliandra* and members of the “Piptadenia group”) in South America and the wider neotropics (Barrett and Parker 2005, 2006; Bontemps et al. 2010; Taulé et al. 2012; Mishra et al. 2012; da Silva et al. 2012; Bournaud et al. 2013, 2017; Silva et al. 2018; Paulitsch et al. 2019a, b, 2020a, b; Mavima et al. 2020), as well as with strains isolated from *Mimosa* growing in pantropical invasive environments.

## 2. Material and Methods

### 2.1. Sampling of plants, nodules and soil

Specimens of *Mimosa* species, including aerial parts (for identification), root nodules, and soil sampled from around the roots were sampled in various sites and altitudes in the states of Bahia (BA), Ceará (CE), Distrito Federal (DF), Espirito Santo (ES), Goiás (GO), Mato Grosso (MT), Minas Gerais (MG), Rio de Janeiro (RJ), Roraima (RR), Santa Catarina (SC) and Sao Paulo (SP) from 2009 to 2018 (Figure 1, Tables 1 and S1). For each survey, branches with leaves were collected (including reproductive structures, when available), and a voucher for each specimen was deposited and then later identified at the herbarium of Embrapa-CENARGEN. In parallel, soil from the rhizosphere was excavated, and nodules (when present) were sampled from the roots visibly connected to the parent plant. The nodules were transferred to sterilized plastic tubes (1 – 2 ml) containing silica gel. Soil samples were also collected, placed in plastic bags, and then dried at room temperature until analysis; some soils were also used for “trapping” rhizobia from *M. pudica* seedlings (see below for details about seed germination).

**Fig. 1.**
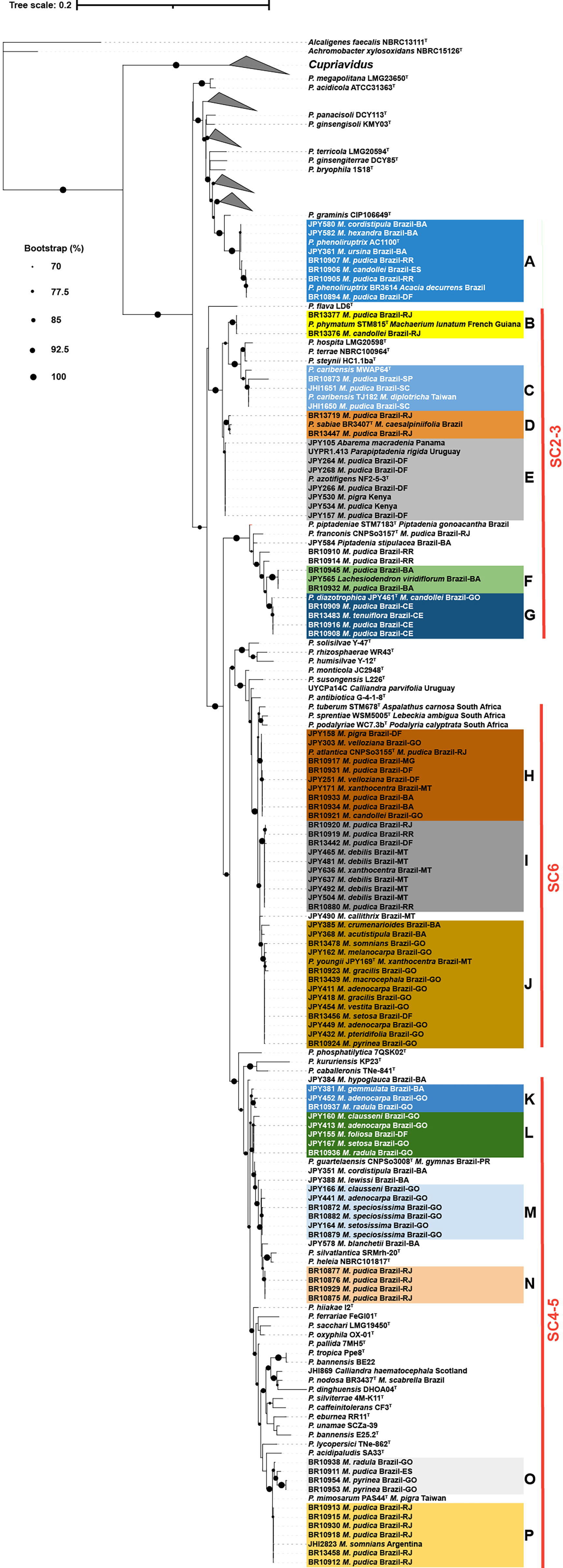
16S rRNA-*recA Paraburkholderia*. Phylogenetic tree built using IQ-TREE, with ultrafast bootstrap analysis (1000 iterations) and the ‘TIM2+F+I+G4’ best-fit model according to Bayesian Information Criterion. The alignment contained 226 concatenated 16S rRNA and *recA* sequences with a total of 1189 nucleotide positions and 250 parsimony-informative sites. The phylogenetic relationships of the genus *Paraburkholderia* are shown and the strains pertaining to the genus *Cupriavidus* were collapsed. Some clades of non-symbiotic *Paraburkholderia* strains were also collapsed to reduce the size of the tree. Phylogenetic clades A to P are color-coded. Plant host species and geographic origin of relevant strains are indicated. For Brazilian strains and when available, the state of origin is also indicated using two-letter abbreviations (*e.g.*, Brazil-DF).

### 2.2. Isolation and screening of bacteria from the nodules

Nodules were washed with sterile deionized water, then surface disinfected by immersion in 95% ethanol for 30 s, followed by immersion in 3% sodium hypochlorite for 5 min, and four washes in sterile distilled water. Nodules were then crushed and streaked onto Medium 79 (Fred and Waksman 1928), otherwise known as yeast mannitol agar or YMA (Vincent, 1970), and incubated at 28°C from 2 to 7 days. Single colonies were transferred to new plates and were then grown in liquid YM media for 2 to 5 days (until visual growth of the bacteria was observed). From each isolate, one 1 ml aliquot was mixed with sterile glycerol 1:1 and frozen at -80°C, while another aliquot was used to extract of DNA, according to Bontemps et al. (2010).

### 2.3. Sequencing of the 16S rRNA, recA, nodC and nifH genes

Novel bacterial strains isolated during the present study (BR-strains, Table S1) were studied by Sanger sequencing of the 16S rRNA gene as well as the housekeeping gene *recA* (recombinase A) and the *nifH* (nitrogenase reductase) and *nodC* (N-acetyl glucosaminyl transferase) symbiosis-essential genes. DNA was extracted from pelleted bacterial cells grown in liquid YM medium using the Wizard Genomic DNA Purification System (Promega). Genes of interest were amplified from template DNA by PCR using GOTaq polymerase (Promega) according to the recommendations of the supplier. Gene-specific cycling conditions were as specified in previous studies, as follows: 16S rRNA genes were amplified using primers 27F and 1492R (Weisburg et al. 1991). To amplify the *recA* gene, different conditions were used for each bacterial genus, as previously described, using the primers Burk recA F and Burk recA R for *Paraburkholderia* described in Mishra et al. (2012), Cupri recA F and Cupri recA R for *Cupriavidus* (Andrus et al. 2012) and 41F and 640R for *Rhizobium* (Vinuesa et al. 2005). The *nodC* gene was amplified using previously described conditions and primers: nodCBurk2 F and nodCBurk2 R for *Paraburkholderia* (Bontemps et al. 2010) and nodCCtai468 F and nodCCtai1231 R for *Cupriavidus* and 540 F and 1160 R for *Rhizobium* (Mishra et al. 2012). Concerning the *nifH* gene, primers used were Burk nifH F and Burk nifH R for *Paraburkholderia* (Chen et al. 2006), nhcf3 and nhcr4 for *Cupriavidus* (Andam et al. 2007) and nifH F and nifH R for *Rhizobium* (Chen at al. 2005a, b); for DNA of challenging strains from any of the three genera, the *nifH* primers PolF and PolR (Poly et al. 2001) were also successfully used. Amplicons were visually inspected by gel electrophoresis and prepared for sequencing by treatment with alkaline phosphatase FastAP and exonuclease ExoI (Thermo Scientific) for 15 min at 37°C followed by 20 min at 85°C. Sequencing reactions were performed bi-directionally using the primers also used for PCR and the Big Dye kit (Applied Biosystem). Purified reaction products were analyzed using a ABI3500 capillary sequencer (Applied Biosystem). Sequence files were quality-processed and assembled into contigs using BioNumerics version 7.1 (Applied Mathematics, Belgium). Sequences were deposited and received Genbank accession numbers (Table S1).

### 2.4. Phylogenetic analysis using the 16S rRNA, recA, nodC and nifH genes

*Alpha-* (*Rhizobium*) and *Beta-* (*Paraburkholderia* and *Cupriavidus*) *proteobacteria* sequences were analyzed separately to prevent long branches and loss of phylogenetic resolution. Multiple sequence alignments of datasets consisting of previously published sequences, the sequences of type strains representative of described species and the sequences obtained in this study were generated in MEGA 7 (Kumar et al. 2016) using MUSCLE (Edgar, 2004). Type strain information was obtained from the List of Prokaryotic names with Standing in Nomenclature (LPSN; https://lpsn.dsmz.de; Parte et al. 2020). The 16S rRNA and *recA* gene datasets were concatenated after aligning and trimming the individual alignments in MEGA 7. Visually inspected multiple alignments were exported in fasta format, and maximum-likelihood phylogenetic trees were constructed using IQ-tree Webserver (Trifinopoulos et al. 2016) with ultrafast bootstrap analysis (Minh et al. 2013) and 1000 iterations. For each alignment, the most appropriate substitution model was selected using the IQ-tree Webserver model selection tool (ModelFinder; Kalyaanamoorthy et al. 2017). Final drawings of phylogenetic trees were generated using iTOL (Interactive Tree of Life, Letunic and Bork 2021).

### 2.5. Genome sequencing and Average Nucleotide Identity (ANI) Analysis

The genomes for reference strains/species were obtained from the National Center for Biotechnology Information (NCBI; Benson et al. 2017, Table S1). The genomes of 15 strains were sequenced during this study (Table S1). The sequencing was performed by MicrobesNG (University of Birmingham, UK) as per Estrada-de los Santos et al. (2018), after which genomes were assembled using SPAdes v3.7 (Bankevich et al. 2012; Nurk et al. 2013). These whole genome assemblies have been deposited in the NCBI database under the accessions listed in Table S1.

These 15 newly-genome-sequenced strains were selected to represent most of the distinct lineages observed within the phylogenies. For all pairs of genome sequences, average nucleotide identity (ANI; using the ANIb algorithm) (Arahal 2014; Goris et al. 2007) values and G+C content ratios were determined using JSpecies (Richter and Rosselló-Móra 2009).

### 2.6. Nodulation Tests

All the BR isolates were tested for their nodulation capability by inoculation of liquid cultures onto *Mimosa pudica* seedlings under sterile conditions. The JPY strains were all confirmed to nodulate their cognate hosts and/or *M. pudica* by Bontemps et al. (2010). For the BR strains, germinated seedlings were obtained by breaking seed dormancy (scarification) with concentrated sulfuric acid for 10 min, followed by surface disinfection (sequential immersion in 97% ethanol for 30 s, 5 min in a 30% hydrogen peroxide solution and finally five washes in sterile water). The seeds were then germinated in Petri dishes with 1% water agar at 28°C for 48 hours in the dark. Germinated seeds were transferred to sterilized glass tubes (200 x 30 mm) containing vermiculite as substrate moistened with a nitrogen-free nutrient solution with pH 6.8 (Norris and T’Mannetje 1964). Each plant was then inoculated with 1 mL of YM broth containing the bacteria of interest in the late logarithmic growth phase; *Cupriavidus taiwanensis* LMG19424_T_ was used as a positive control (Elliott et al. 2007a) and sterile culture medium as a negative control. Plants (n=3 per treatment) were grown in a growth room for 45 days at 28°C and with a 16 h light photoperiod. Bacteria were re-isolated from any nodules formed, and their identity confirmed by genomic profile comparison to the inoculated strains by BOX-PCR (Koeuth et al. 1995).

### 2.7. Soil analysis

Soil analysis was conducted in the Agricultural Chemistry Laboratory of Embrapa Agrobiologia using procedures described in Silva et al. (2009). Briefly, the soil was first air dried and sieved to 2 mm, removing plant material. The pH of the sampled soil in water was then measured using a pH-electrode, while Ca^2+^, Mg^2+^, and Al^3+^ were extracted using 1 M KCl and analyzed by atomic absorption spectroscopy. Phosphorus (P) and potassium (K) were extracted using the Mehlich I solution (Silva et al. 2009), and P was analyzed by UV-spectrophotometry, and K by atomic absorption.

### 2.8. Statistical analyses

Analyses were performed for those rhizobial isolates from widespread *Mimosa* species for which soil samples were also taken (mainly *M. pudica*; Table S1), in order to identify correlations between biological (occurrence of bacterial genera *Cupriavidus*, *Paraburkholderia* and *Rhizobium*) and soil chemistry attributes (pH, Al+3, Ca, Mg, P and K), a multivariate redundancy analysis (RDA) was performed in R (R Core Team, 2021). The Monte Carlo permutation test assessed the significance of relationships between biological and chemical data matrices, considering 5% as the probability of significance. The soil chemistry variables were centralized and standardized. The heatmap was constructed using the pheatmap package in R and rows were clustered using hclust based on Eucledian distance.

## 3. Results

### 3.1. Nodulation of Mimosa species in eleven states of Brazil and diversity of their symbiotic bacteria

Nodules from 10 *Mimosa* taxa were sampled in 11 Brazilian states, with *M. pudica* sampled from all states (Table S1, Fig. S1). Of these, *M. macrocephala* from the Cerrado biome in GO was a new report of nodulation. These, together with data from the recent studies of Paulitsch et al. (2019a, b) and Klepa et al. (2021) raises the total of known nodulated *Mimosa* species to 135 when they are added to those published in earlier studies (Sprent 2009; Lammel et al. 2013, 2015; Bontemps et al. 2016; Platero et al. 2016; Pires et al. 2018).

We obtained 79 potentially symbiotic isolates from the nodules of these *Mimosa* spp., including 60 isolates from *M. pudica* nodules (Table S1). Phylogenies of these new isolates (all prefixed with “BR”) were then constructed using their 16S rRNA sequences, to which were added sequences from 35 strains (all prefixed with “JPY”) described in a previous study of *Mimosa* symbionts from central Brazil (Bontemps et al. 2010) together with reference/type strains plus some unpublished strains from the authors’ strain collections (Table S1). Based on 16S rRNA gene partial sequences most isolates were Beta-rhizobia, with the majority belonging to *Paraburkholderia*, plus a few belonging to *Cupriavidus* and *Rhizobium* (Fig. S1). However, all 35 JPY strains were *Paraburkholderia,* confirming their former affiliation to the genus *Burkholderia*, the generic name under which they were originally published (Bontemps et al. 2010).

The *recA* sequences of 77 of the BR isolates were also obtained (Fig. S2); these were combined with those of the 35 JPY strains to construct a 16S rRNA-*recA* phylogeny (Figure 1), which allows for greater resolution in terms of which Beta-rhizobial species the BR and JPY isolates belonged to. Although many of the JPY strains from the study of Bontemps et al. (2010) have already been allocated to various species, such as *P. mimosarum* (Chen et al. 2006), *P. nodosa* (Chen et al. 2007), *P. sabiae* (Chen et al. 2008), *Trinickia* (syn. *Burkholderia*) *symbiotica* (Sheu et al. 2012), *P. diazotrophica* (Sheu et al. 2013), *P. atlantica* and *P. youngii* (Mavima et al. 2020), the 35 JPY strains re-examined here were originally placed in six of the seven 16S rRNA-*recA* SC but with no apparent affiliation to any described species at the time that they were first published (Bontemps et al. 2010). Here, we more firmly establish the identity of these so far unclassified JPY *Paraburkholderia* strains and determine the identities of the new BR strains *vis-à-vis* current revisions in the taxonomy of Beta-rhizobia. A concatenated 16S rRNA-*recA* phylogeny reveals that the combined BR-JPY isolates belong to 18 genotypes of beta-rhizobia comprising 16 *Paraburkholderia* (Fig. 1, Table S1) and two *Cupriavidus* genotypes (Fig. 2, Table S1), plus a single genotype of *Rhizobium*.

**Fig. 2.**
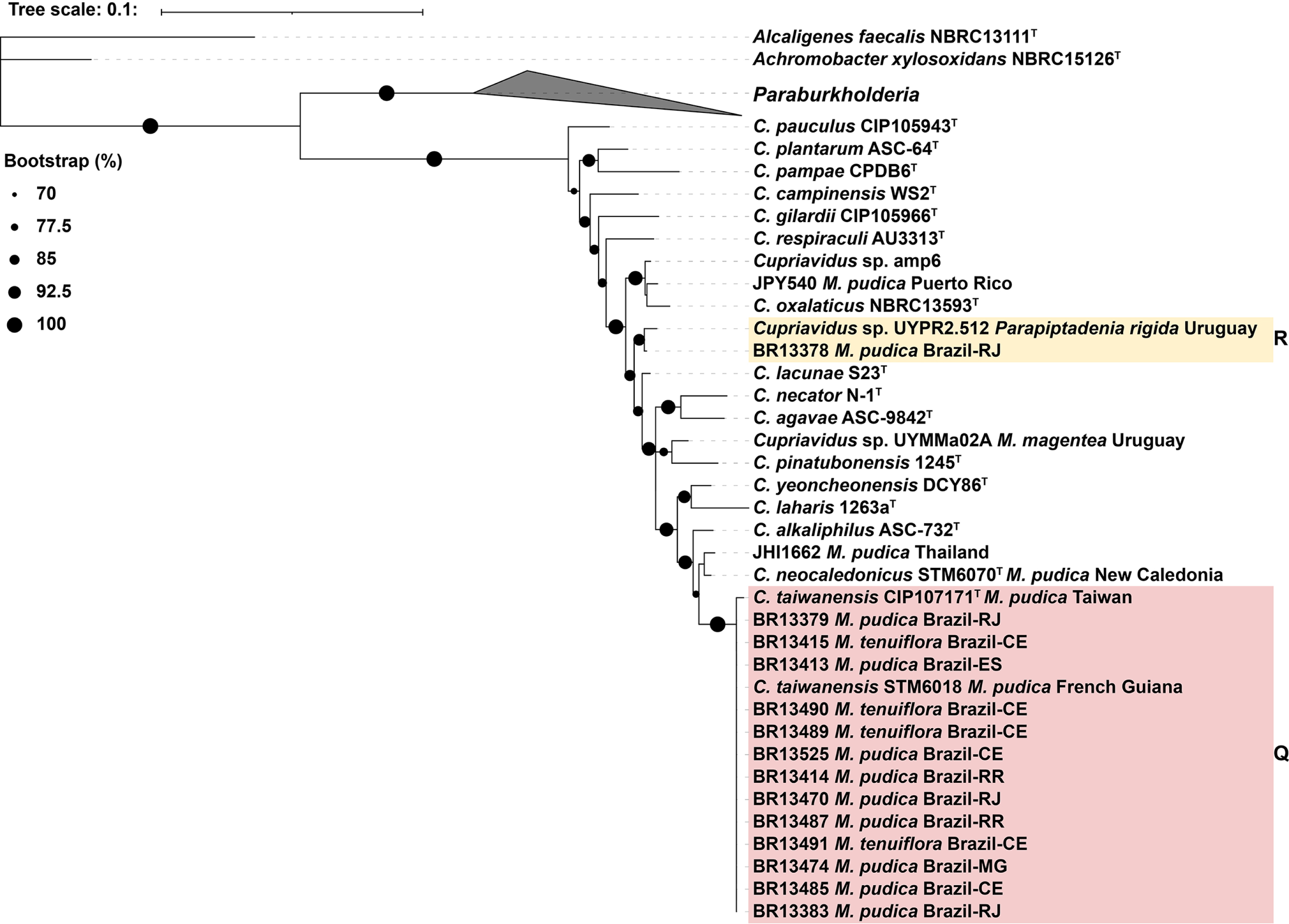
16S rRNA-*recA Cupriavidus*. Phylogenetic tree built using IQ-TREE, with ultrafast bootstrap analysis (1000 iterations) and the ‘TIM2+F+I+G4’ best-fit model according to Bayesian Information Criterion. The alignment contained 226 concatenated 16S rRNA and *recA* sequences with a total of 1189 nucleotide positions and 250 parsimony-informative sites.. The phylogenetic relationships of the genus *Cupriavidus* are shown and the genus *Paraburkholderia* was collapsed. Plant host species and geographic origin of relevant strains are indicated. For Brazilian strains and when available, the state of origin is also indicated using two-letter abbreviations (*e.g.*, Brazil-DF).

Nine of the rhizobial genotypes clustered with type strains from validly described species of *Paraburkholderia*, such as: *P. phenoliruptrix* (**Group A** *Mimosa*-associated strains from BA, DF, ES and RR plus the *Acacia decurrens* symbiont, BR3614); *P. phymatum* (**Group B** *Mimosa* symbionts from RJ); *P. caribensis* (**Group C** *Mimosa*-associated strains from SC and SP); *P. sabiae* (**Group D** *Mimosa* symbionts from RJ); *P. azotifigens* (**Group E** *Mimosa*-associated strains from DF and Kenya, plus *Parapiptadenia* and *Jupunba*/*Abarema* symbionts from Uruguay and Panama, respectively); *P. diazotrophica* (**Group G** *Mimosa*-associated strains from CE); *P. atlantica* and *P. youngii* (**Groups H** and **J,** respectively; *Mimosa*-associated strains from BA and central Brazil: DF, GO, MT), and *P. mimosarum* (**Group P** *M. pudica* symbionts from RJ and Argentina).

The remaining groups in the *Paraburkholderia* 16S rRNA-*recA* phylogeny did not contain any type strains. These were: **Group F** (closest to *P. diazotrophica*) comprising *Mimosa* symbionts from BA and RR, and JPY565 from the “Piptadenia group” species *Lachesiodendron viridiflorum* (syn. *Piptadenia viridiflora*) in BA; **Group I** (closest to *P. atlantica/P. youngii*) comprising *Mimosa* symbionts from DF, MT, RJ and RR; the related **Groups K, L and M** (closest to *P. guartelaensis*) comprising *Mimosa*-associated strains from BA, DF and GO; **Group N** (closest to the non-symbionts *P. silvatlantica* and *P. heleia*) comprising *Mimosa* symbionts from GO and RJ; and **Group O** (closest to *P. mimosarum*) comprising *Mimosa*-associated strains from GO and ES. There were also ten single strain lineages based upon the approach of recognizing **Groups** on the basis of the smallest recovered, supported monophyletic lineage: JPY351, JPY388, JPY384 and JPY578 from endemic *Mimosa* spp. in BA that were all close to *P. guartelaensis* and **Groups M and N**; JHI869 isolated from nodules on *Calliandra haematocephala* at the Royal Botanical Gardens Edinburgh, UK (this study), which was close to *P. nodosa*; UYCPa14C isolated from *Calliandra parvifolia* in Uruguay (Langleib et al. 2019), which was closest to the non-symbiont *P. antibiotica*; JPY490 from the MT endemic *M. callithrix* which was nested between *P. atlantica* and *P. youngii* but belonged to neither species; and JPY584 isolated from *Piptadenia stipulacea* in BA (Bournaud et al. 2013) plus BR10910 and BR10914 from *M. pudica* in RJ, which were both closest to *P. franconis*, *P. diazotrophica* and to **Group F**.

The 16S rRNA-*recA* phylogeny suggested at least six potential new species among the BR-JPY *Paraburkholderia* strains. To further investigate this, we sequenced the whole genomes of representative strains from **Groups F, I, K, L, and M** (strains from **Groups N** and **O** were unavailable for genome sequencing at this time) and performed an ANI analysis on them compared with the closest type strains in the 16S rRNA-*recA* phylogeny. This analysis (Table S2a) demonstrated that the **Group F** strain JPY565 from *L. viridiflorum* (Bournaud et al. 2013) shared low similarities (<93%) with all type strains in the *P. diazotrophica-P. franconis-P. piptadeniae* Species Complex or SC (equivalent to SC2-3 of Bontemps et al. 2010), the **Group I** strain JPY481 isolated from *M. debilis* in the Pantanal wetlands of MT shared slightly more than 94% similarity with *P. atlantica* and *P. youngii*, while three strains from GO in the *P. mimosarum*-*P. nodosa*-*P. guartelaensis* SC (SC4-5 of Bontemps et al. 2010), JPY452 (**Group K,** *M. adenocarpa*), JPY167 (**Group L**, *M. setosa*), and JPY164 (**Group M**, *M. setosissima*), shared <94% similarity with the type strains of *P. nodosa* and *P. guartelaensis,* but are related to each other, *e.g.*, with JPY164 and JPY167 sharing >98% similarity. Strain JHI869 isolated from *Calliandra haemotocephala* nodules at the Royal Botanic Gardens Edinburgh (RBGE), Edinburgh, UK, was a singleton in the 16S rRNA-*recA* phylogeny, and its genome shared only 90% with the closest type strain, *P. nodosa*, while the genome of JHI2823, which was isolated from the widespread neotropical species *M. somnians* sampled in northern Argentina, shared >98% similarity with *P. mimosarum*. Finally, the genome of JPY584 from *Piptadenia stipulacea* in BA (Bournaud et al. 2013) shared <93% similarity with *P. diazotrophica, P. franconis* and *P. piptadeniae*. The ANI thus suggests that **Group F**, **Group I**, **Group K**, and **Groups L+M**, may represent four new *Paraburkholderia* species.

The 16S rRNA-*recA* phylogeny also suggested that the nodulating strains in **Group E** belonged to *P. azotifigens*. This species was isolated from paddy field soil in South Korea (Choi and Im 2018) and was previously considered to be a free-living diazotroph with no capacity to nodulate legumes. It was tested by comparing the genomes of the **Group E** strains JPY530 and JPY534 (isolated from *M. pigra* and *M. pudica,* respectively, in Kenya), JPY105 (AMAC11-3) (isolated from the mimosoid tribe Ingeae species *Jupunba macradenia* in Panama; Barrett & Parker, 2005), and UYPR1.413 (isolated from the “Piptadenia group” species *Parapiptadenia rigida* in Uruguay; Taulé et al. 2012; De Meyer et al. 2015b) with that of the type strain of *P. azotifigens*, NF5-2-3^T^, the comparison showed these genomes shared more than 98% similarity (Fig. S4).

Of the 14 *Cupriavidus* strains isolated in this study (prefixed BR), most were from nodules on the widespread species *M. pudica* and *M. tenuiflora* sampled in CE, ES, MG, RJ and RR, and all but one was placed in **Group Q** with *C. taiwanensis* in the 16S rRNA-*recA* phylogeny (Fig. 2). The exception was BR13378 from *M. pudica* in RJ, which was placed in **Group R** together with UYPR2.512 from *Parapiptadenia rigida* nodules in Uruguay (Taulé et al. 2012; De Meyer et al. 2015a); the type strain of *C. lacunae* was closest to this pair of strains. Other divergent strains were UYMMa02A from *M. magentea* in Uruguay (Platero et al. 2016; Iriarte et al. 2016), which was closest to *C. pinatubonensis*, JPY540 and amp6 from *M. pudica* in Puerto Rico and Texas, respectively, which were closest to each other and to *C. oxalaticus*, and JHI1662 from *M. pudica* nodules sampled in northern Thailand, which was related to *C. neocaledonicus*, a symbiont of *M. pudica* in New Caledonia (Klonowska et al. 2020). The genomes of both JPY540 and JHI1662 were sequenced for this study so their relatedness to these type strains could be tested using ANI (Table S2b); this revealed that JPY540 shared <91% similarity with *C. oxalaticus*, while JHI1662 shared <93% similarity with *C. neocaledonicus*.

The final genotype (**Group S**) was *Rhizobium* (Fig. S3). These fourteen strains were all isolated from *M. pudica* in CE, DF, MG, and RR. The ten DF strains were previously described as belonging to the new species *R. altiplani* (Barauna et al. 2016). Of the remaining four strains, BR13443 was isolated from seedlings grown in soil from MG, and it clustered with *R. pisi* and *R. etli*, while strains BR10892 (CE) and BR13410 (RJ) were grouped with *R. tropici* and *R. multihospitium*; the two strains from RR, BR10891 and BR10904, were close to *R. mesoamericanum*.

### 3.2. Symbiosis-essential genes

Two symbiosis-related genes, *nodC* and *nifH*, were examined in this study (Fig. 3, 4). For the *Paraburkholderia* strains, both genes were generally congruent with each other and with the 16S rRNA-*recA* phylogeny, forming groups that coincided with each of the three largest SC identified by Bontemps et al. (2010) for *Mimosa*-nodulating (*Para)burkholderia i.e.*, SC2-3 comprising *P. phymatum*-*P. sabiae*-*P. diazotrophica*-*P. caribensis*-*P. azotifigens* (**Groups B – G** in the present study), SC4-5 comprising *P. mimosarum*-*P. nodosa*-*P. guartelaensis* (**Groups K – P** in the present study), and SC6 comprising the ex-*P. tuberum* sv. mimosae species *P. atlantica* and *P. youngii* (**Groups H – J** in the present study).

**Fig. 3.**
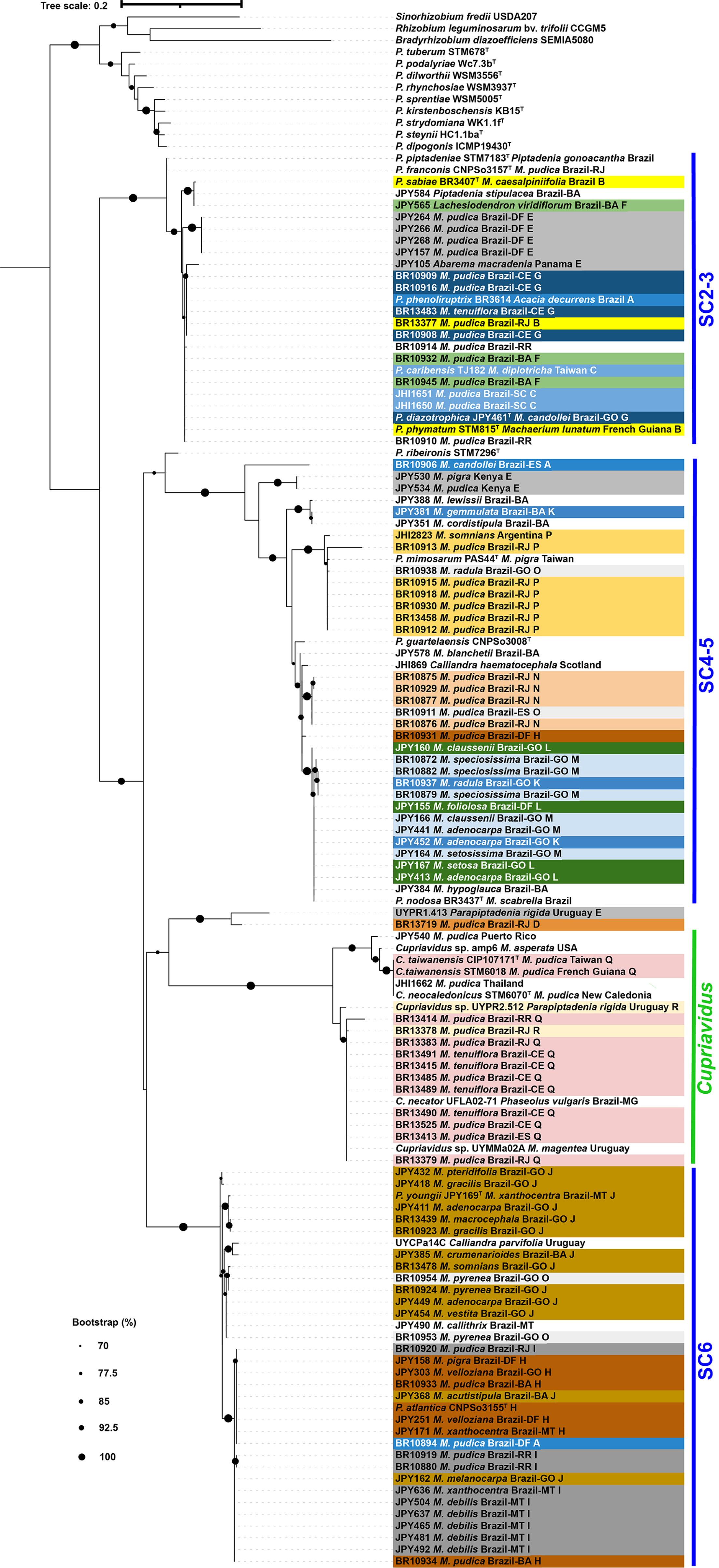
nodC. Phylogenetic tree built using IQ-TREE, with ultrafast bootstrap analysis (1000 iterations) and the ‘HKY+F+I+G4’ best-fit model according to Bayesian Information Criterion. The alignment contained 132 sequences, 318 nucleotide positions and 156 parsimony-informative sites. Plant host species and geographic origin of relevant strains are indicated. For Brazilian strains and when available, the state of origin is also indicated using two-letter abbreviations (*e.g.*, Brazil-DF).

Some strains showed incongruency in that they were placed in a *nodC* **Group** that was different from their 16S rRNA-recA SC/**Group**. These included two *P. mimosarum*-like strains from the GO endemic *M. pyrenea*, BR10953 and BR10954 (both 16S rRNA-*recA* SC4-5/**Group O**), that were all nested in a *nodC* lineage that corresponds to 16S rRNA-*recA* SC6/**Groups H-J** (*P. atlantica*-*P. youngii*); the *P. sabiae*-like strain BR13719 (16S rRNA-*recA* SC2-3/**Group D**) isolated from *M. pudica* in RJ whose *nodC* sequence was distinct from all the SC/**Groups**, being most closely related to UYPR1.413 isolated from *Parapiptadenia rigida* in Uruguay (Taulé et al. 2012). Moreover, the *P. azotifigens*-like strains JPY530 and JPY534 (16S rRNA-*recA* SC2-3/**Group E**) from Kenya, were divergent from all the SC/**Groups** in their *nodC* phylogenies, while the *nodC* sequence of the *P. atlantica* strain BR10931 from *M. pudica* in DF (16S rRNA-*recA* SC6/**Group H**) was divergent, instead grouping closer to *P. nodosa* (16S rRNA-*recA* SC4-5). The *Paraburkholderia nifH* phylogeny (Fig. 4) gave a similar picture to the *nodC* phylogeny (Fig. 3), but there were some minor differences.

**Fig. 4.**
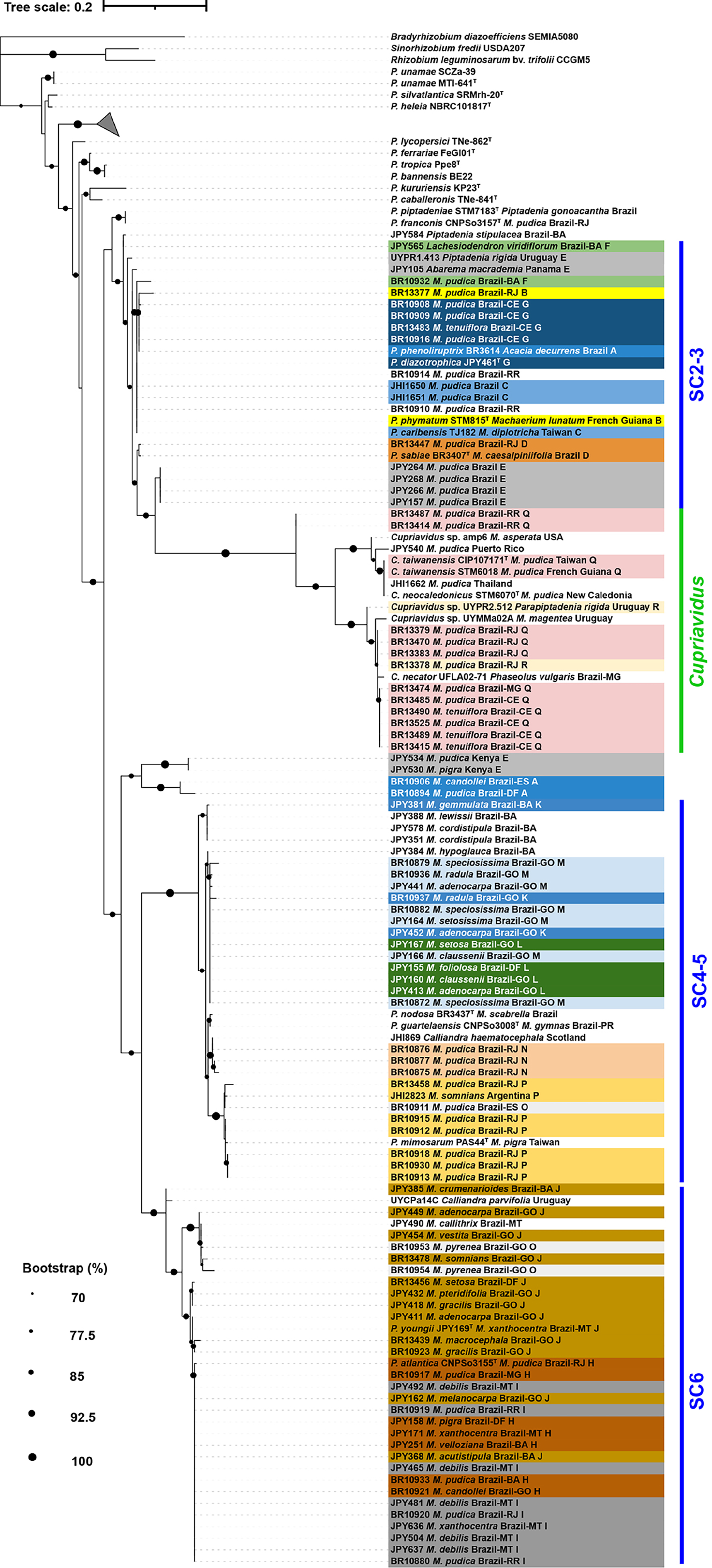
nifH. Phylogenetic tree built using IQ-TREE, with ultrafast bootstrap analysis (1000 iterations) and the ‘TN+F+I+G4’ best-fit model according to Bayesian Information Criterion. The alignment contained 141 sequences, 246 nucleotide positions and 99 parsimony-informative sites. Plant host species and geographic origin of relevant strains are indicated. For Brazilian strains and when available, the state of origin is also indicated using two-letter abbreviations (*e.g.*, Brazil-DF). The collapsed clade represented by a grey triangle contains *P. xenovorans* LB400^T^, *P. dioscoreae* Msb3^T^, *P. aromaticivorans* BN5^T^, *P. strydomiana* WK1.1f^T^, *P. podalyriae* WC7.3b^T^, *P. sprentiae* WSM5005^T^, *P. tuberum* STM 678^T^, *P. steynii* HC1.1ba^T^, *P. dipogonis* ICMP19430^T^, *P. kirstenboschensis* Kb15^T^, *P. dilworthii* WSM3556^T^, *R. rhynchosiae* LMG27174^T^

Particular mention should be made here about the symbiotic strains in **Group A**, which are closely-related or belonging to *P. phenoliruptrix*, which was originally described as a non-symbiont, and is closer to non-symbionts in the 16S rRNA-*recA* phylogeny than any of the other *Mimosa*-nodulating *Paraburkholderia* groups (Fig. 1). One **Group A** strain from the introduced Australian legume *Acacia decurrens*, BR3614, possessed *nodC* and *nifH* genes that placed them in SC2-3 (Fig. 3, 4), but two other strains, the *M. pudica* symbiont BR10894 from DF and strain BR10906, which was isolated from *M. candollei* in ES, were placed either in SC6 (BR10894) or in a separate lineage close to SC4-5 (BR10906) in the nodC phylogeny. Moreover, in the nifH phylogeny both these Brazilian strains grouped with a pair of Kenyan strains, JPY530 and JPY534, forming two diverged paired *nifH* lineages, neither of which could be placed in *nifH* lineages that correspond to any of the 16S rRNA-*recA* SC/**Groups**. Strain BR10894 was, therefore, incongruent across all three phylogenies (16S rRNA-*recA*, *nodC* and *nifH*).

The *nodC* and *nifH* sequences of all but one of the BR *Cupriavidus* strains were grouped with a *Mimosa*-nodulating *C. necator* strain (UFLA02-71) isolated from nodules of common bean (*Phaseolus vulgaris*) and *Leucaena leucocephala* grown in soil sampled from a pasture in MG, Brazil (Silva et al. 2012). It was a substantial incongruency given that all the BR strains (except for the *C. lacunae*-like BR13378) were placed in *C. taiwanensis* according to the 16S rRNA-*recA* phylogeny, including BR13474 (Fig. 2) that was isolated from *M. pudica* growing in the same field from which the soil that was used to trap UFLA02-71 was sampled (James and de Faria, unpublished). Strain UYMMa02A from *M. magentea* in Uruguay and the *C. lacunae*-like BR13378 were also clustered in this *nodC*/*nifH* group, but UYPR2.512 from *P. rigida* (Taulé et al. 2012; de Meyer et al. 2015a) was slightly divergent from it. The *nodC* and *nifH* sequences of the *C. oxalaticus*-like strains, JPY540 and amp6, isolated, respectively, from *M. pudica* in Puerto Rico (this study) and *M. asperata* in Texas (Andam et al. 2007; De Meyer et al. 2015c), were both slightly divergent from *C. taiwanensis*. The only *nifH*-specific incongruency in the *Cupriavidus* collection was that of two *C. taiwanensis* strains isolated from *M. pudica* in RR (BR13414, BR13487); these were paired together and were substantially divergent from all the other strains.

### 3.3. Effects of host species, geography and edaphic factors on rhizobial diversity

There were some potential relationships evident between the 16S rRNA-*recA* Groups and their (mainly) *Mimosa* host plants (Fig. 5). For example, **Group N** comprises only four strains isolated from *M. pudica* sampled in RJ and GO, and **Group P** comprises only *M. pudica* strains from RJ. On the other hand, although *M. pudica* strains were present in 12 out of the 16 *Paraburkholderia* Groups (**Groups A – I, N – P**), there were none in **Groups J – M**; these Groups only contained strains (including *P. youngii*) mainly isolated from *Mimosa* spp. endemic to the highlands of GO, MT and BA (Simon and Proença 2000; dos Reis Junior et al. 2010). As mentioned previously, *Cupriavidus* (**Groups Q and R**) and *Rhizobium* (**Group S**) were only associated with the widespread species *M. pudica*, although **Group Q** was also isolated to a lesser extent from *M. tenuiflora*, another widespread species.

**Fig. 5.**
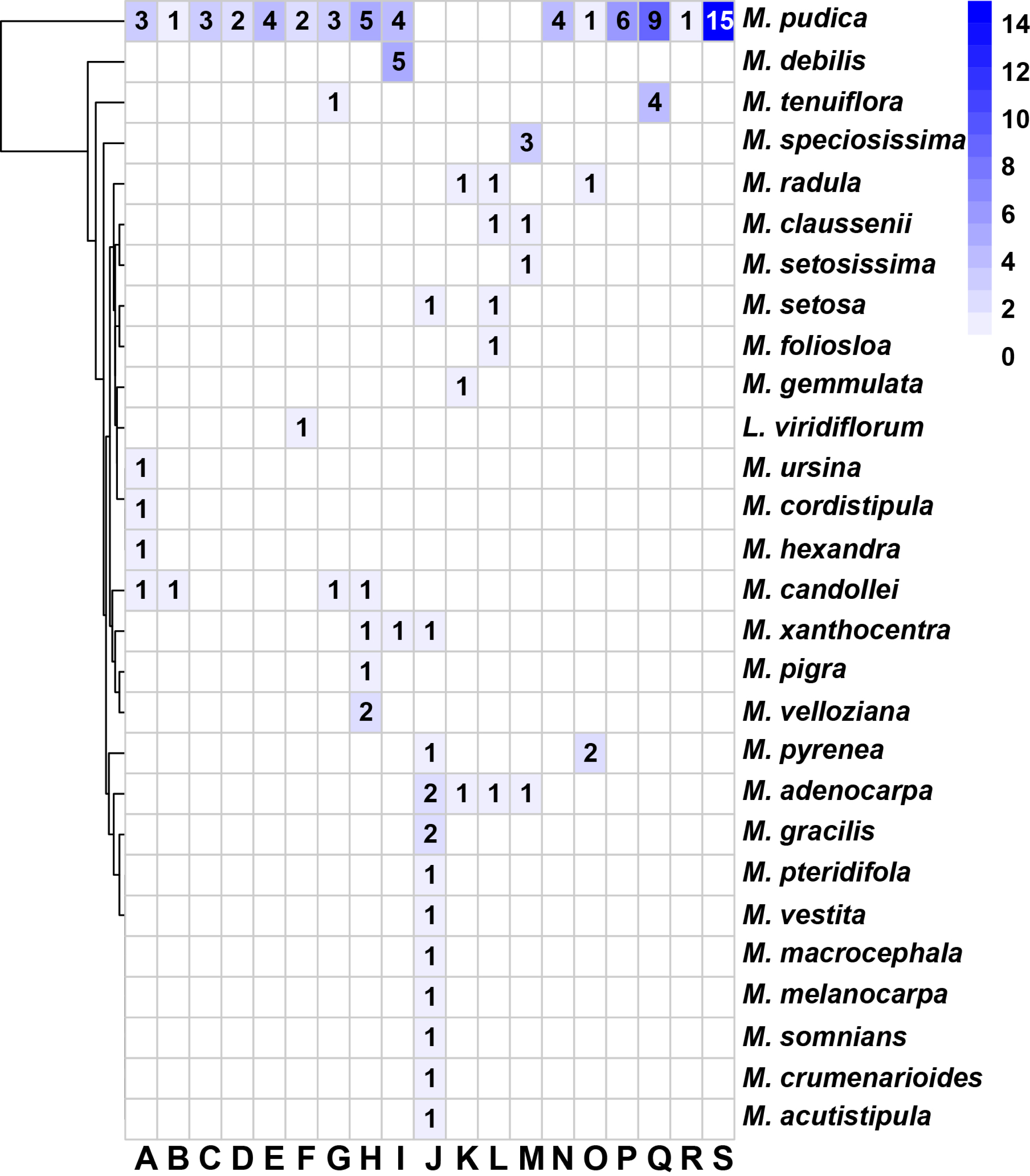
Heatmap showing the relationship of phylogenetic clades with host-plant origin. Absolute numbers of strains are indicated in heatmap cells. Clade numbers are indicated under the columns.

The geographical distribution of the 16S rRNA-*recA* Groups was challenging to determine with any precision, as there were substantial differences in sampling, and there was also some sampling bias, especially in states where particular locations harboring *M. pudica* were more densely sampled (*e.g.*, *Rhizobium altiplani* in DF; Barauna et al. 2016) (Fig. 6 map). However, if we look at the states in which many endemic *Mimosa* spp. were sampled, such as BA and GO, it can be seen that the dominant 16S rRNA-*recA* **Groups** were those mostly associated with endemic *Mimosa* spp., *e.g.*, *Paraburkholderia* **Groups J – M**, **and O**, but also in MT in which *Paraburkholderia* **Group I** was consistently isolated from native/endemic *Mimosa* spp. These states thus highlight the tight links between the endemic plants, their location, and their microsymbionts. On the other hand, in another of the more densely sampled states, RJ, but which harbors no endemics, there was also a wide diversity of 16S rRNA-*recA* genotypes, albeit ones more commonly isolated throughout the neotropics and within the invasive pantropical range of *M. pudica*; these were *Cupriavidus* (**Group Q and R**), *P. mimosarum* (**Group P**), *P. phymatum*, and *Rhizobium* (**Group S**). Contrastingly, however, RJ was also the only Brazilian state that yielded *Paraburkholderia* **Group N** strains (all isolated from *M. pudica*).

**Fig. 6.**
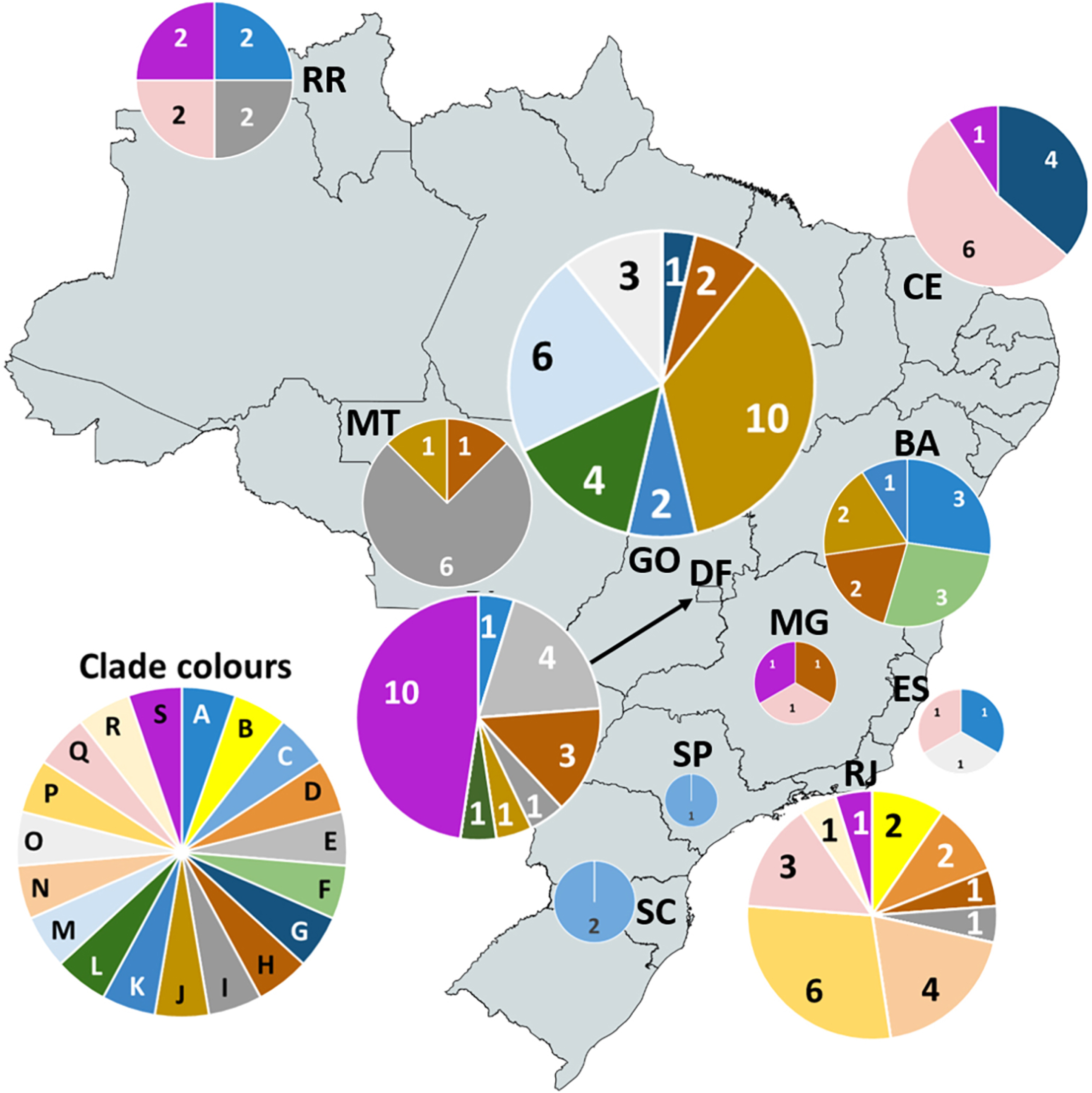
Geographic distribution of strains from different phylogenetic clades in Brazil. Brazilian states are indicated by two-letter codes: BA (Bahia), CE (Ceara), DF (Distrito Federal), ES (Espirito Santo), GO (Goias), MG (Minas Gerais), MT (Mato Gross), SJ (Rio de Janeiro), RR (Roraima), SC (Santa Catarina), SP (Sao Paulo). Pie graphs show the number of strains for each clade per state. Clade colors as in the 16S rRNA-*recA*, *nodC* and *nifH* phylogenies (**Figs 1, 2, 3 and 4**).

The effect of soil characteristics on the diversity and distribution of symbionts of *M. pudica* and other widespread species was tested via an RDA. It used the 62 nodule isolates for which soil data were available from the sites where their hosts were excavated (Table S1). Of the total variation occurring in the biological data, the first two axes (RDA1 and RDA2) could explain 37.80% of the total variation, whereas the remainder (62.2%) corresponded to residual variance (Table 1 and Fig. 7). It can be seen in Fig. 7 that the soil variables pH and Ca are located on the right of the first ordering axis (RDA 1) together with the genus *Rhizobium,* indicating that the occurrence of *Rhizobium* correlated positively with soil pH and Ca. On the negative side of RDA 1, the variable Al^3+^ positively correlated with the genus *Paraburkholderia* but negatively correlated with soil pH and Ca. *Paraburkholderia* did not correlate with P, K and Mg. Concerning RDA 2, *Cupriavidus* was positively correlated with K, Mg and P. Therefore, in soils with higher pH and Ca values, there is a lower chance of finding *Paraburkholderia* and a higher chance of finding *Rhizobium*. More acidic and aluminum rich soils tend to favor *Paraburkholderia,* while mildly acidic soils with higher K, Mg and P tend to favor the genus *Cupriavidus*.

**Fig. 7.**
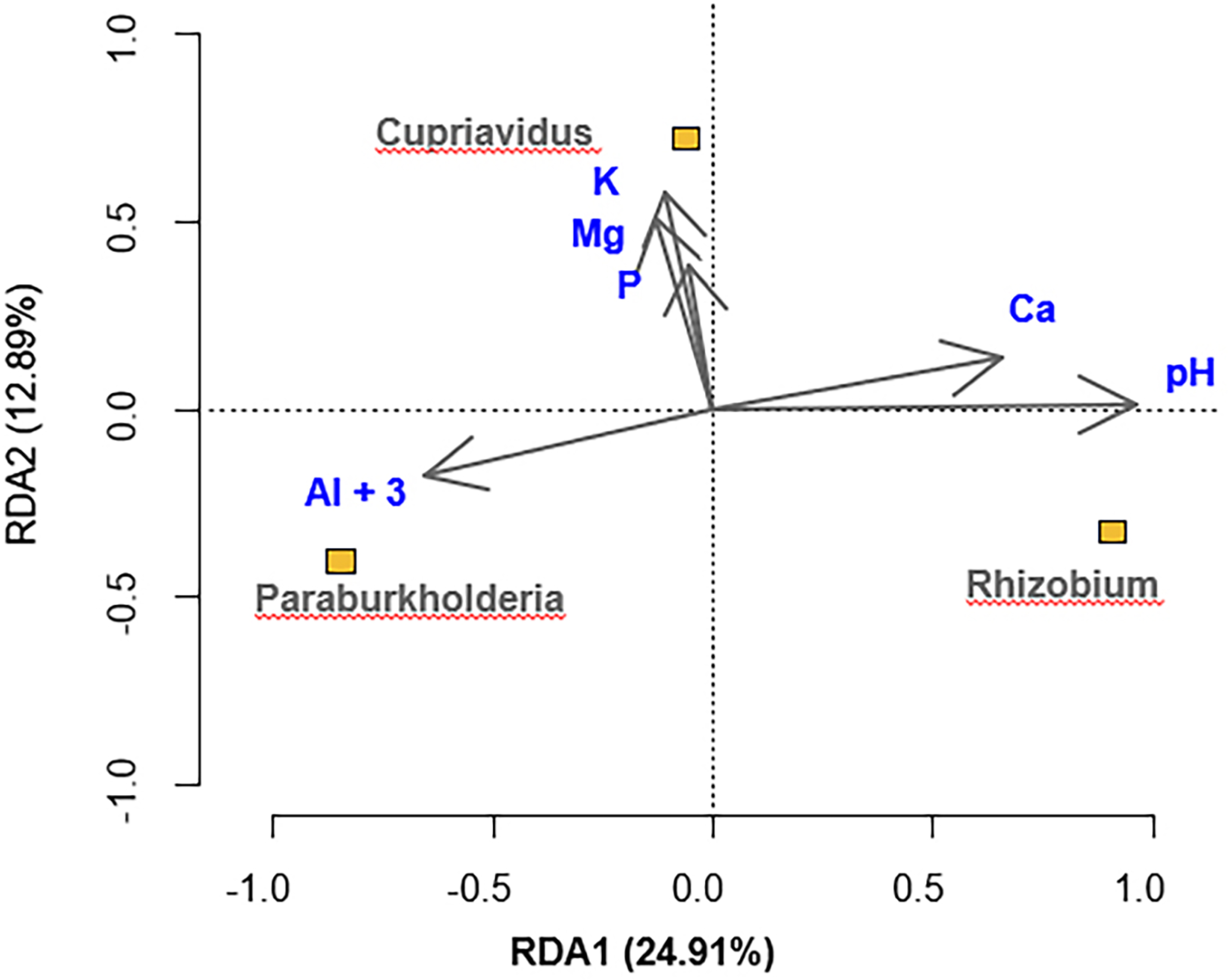
Ordination plot for the first two axes of a (RDA_1_ e RDA_2_) resulting from a redundancy analysis (RDA) between microbiological and soil chemical variables. The soil chemistry data are potentially related to the distribution of the bacterial genera *Paraburkholderia*, *Cupriavidus* and *Rhizobium* that were isolated as nodulating symbionts of *Mimosa* spp. growing in Brazilian soils. Canonical axes are significant according to the Monte Carlo permutation test (F=23.236; P=0.001, 999 permutations).

**Table 1.** Redundancy Analysis (RDA) for soil chemistry data potentially related to the distribution of the bacterial genera *Paraburkholderia*, *Cupriavidus* and *Rhizobium* that were isolated as nodulating symbionts of *Mimosa* spp. growing in Brazilian soils. Canonical axes are significant according to the Monte Carlo permutation test (F=23.236; P=0.001, 999 permutations).

| Chemical variable | RDA1 | RDA2 |
| --- | --- | --- |
| pH | 0.96 | 0.01 |
| Al+3 | -0.65 | -0.18 |
| Mg | -0.13 | 0.51 |
| Ca | 0.65 | 0.13 |
| P | -0.05 | 0.38 |
| K | -0.11 | 0.57 |
| Autovalues | 0.2491 | 0.1289 |
| Cumulative % of explained variance: |  |  |
| Variables of soil bacterial genera | 24.91 | 37.80 |
| Variables of soil bacterial genera and chemistry data | 65.9 | 100.0 |
| Sum of all canonical autovalues (%) |  | 37.80 |

## 4. Discussion

### 4.1. A high diversity of Paraburkholderia nodulating Mimosa in Brazil is contained within three large Species Complexes

In the present study, we took a different approach towards uncovering the diversity of symbionts of the speciose genus *Mimosa* by not focusing so much on the enormous variety of endemic Brazilian species but rather on the symbionts of widespread species that have become pantropical weeds because of centuries of human-mediated dispersal, particularly *M. pudica*. We then combined the dataset from the present study with that of the earlier study of Bontemps et al. (2010) on mainly endemic *Mimosa* spp. to establish if and how the symbionts of these two groups of *Mimosa* differ and if they interact with each other’s hosts. Surprisingly, given that it is now more than 15 years since their study was conducted, the Species Complexes (SCs) demonstrated by Bontemps et al. (2010) for *Mimosa*-nodulating *Paraburkholderia* were confirmed by this new combined dataset. No novel SCs were revealed either in this study or in others published since 2010, with only the species *P. ribeironis*, which was isolated from *Piptadenia gonoacantha* (Bournaud et al. 2013, 2017), being divergent from the already known SCs. On the other hand, the various species that comprise the SCs of Bontemps et al. (2010) are now being progressively described. These include *P. diazotrophica* (SC2-3), *P. franconis* (SC2-3), *P. guartelaensis* (SC4-5), and the ex-“*B. tuberum* sv. mimosae” species *P. atlantica* and *P. youngii* (SC6) (Sheu et al. 2013; Paulitsch et al. 2019a, b, 2020a; Mavima et al. 2021, 2022). In addition, based on groups without any type strains closely affiliated to them, the combined 16S rRNA-*recA* dataset from the present study suggested that there might be at least another six new species (not including single strain lineages). This was reinforced by the ANI analysis using full genomes of representative strains from four of the unaffiliated 16S rRNA-*recA* groups, *i.e.*, Groups F, I, K, and L+M, representing all three of the large SCs. The other two potentially new species suggested by the 16S rRNA-*recA* phylogeny were based on Groups N and O (both SC4-5), but further progress with these awaits a reference strain from each group having its full genome sequenced.

The other important discovery arising from our combined dataset is that the “minor” SC2 and SC3 of Bontemps et al. (2010), which we have combined into SC2-3, is now revealed to be at least as large as the two “major” SCs, SC4-5 and SC6. The expansion of SC2-3 from that first reported by Bontemps et al. (2010) is a consequence of including so many symbionts of widespread lowland *Mimosa* species that were not investigated by this earlier study of mainly endemic species. SC2-3 now comprises *P. phymatum*, *P. sabiae*, *P. diazotrophica*, *P. franconis*, and *P. piptadeniae*, all of which are frequently isolated from *M. pudica* and other widespread *Mimosa* spp. in Brazil and elsewhere in the tropics, but also from non-*Mimosa* mimosoids like *Piptadenia*, *Lachesiodendron*, *Parapiptadenia* and *Jupunba* (*Abarema*) in Brazil (Bournaud et al. 2013), Uruguay (Taulé et al. 2012), and Panama (Barrett and Parker 2005), confirming the cosmopolitan nature of this SC.

Another striking aspect of the newly expanded SC2-3 was the apparent abundance of two previously described non-symbiotic and non-diazotrophic species as *Mimosa*-nodulating bacteria, *i.e.*, *P. azotifigens* and *P. caribensis*; these two species join with *P. phenoliruptrix*, which is in a clade divergent from all those containing *Mimosa* symbionts. *Paraburkholderia caribensis* was first reported to be a symbiont of *Mimosa* in Taiwan by Chen et al. (2003a), but the (non-symbiotic) type strain MWAP64^T^ was isolated originally from soils in Martinique (Achouak et al. 1999); since then, *P. caribensis* has been isolated from nodules of invasive *Mimosa* species in various locations in Asia, such as China (Liu et al. 2020), but this is the first report of it as a symbiont of *Mimosa* in Brazil. Similarly, *P. phenoliruptrix* was originally isolated from a chemostat (Coenye et al. 2004) and then revealed to be related to a symbiont of *M. flocculosa* in Brazil, as represented by *Burkholderia* strain BR3462 (Chen et al. 2005a), which was later formally described as *P. phenoliruptrix* strain BR3459a (Cunha et al. 2012), and which like BR3614 (this study), possesses *P. phymatum*-type *nod* and *nif* genes (Soares-Neto et al. 2022). Thereafter, *P. phenoliruptrix* has been quite frequently isolated as a nodulating symbiont of native *Mimosa* and *Calliandra* species in Brazil, and with widely varying nod and nif genes (this study), but also of introduced species like *Acacia decurrens* (Zilli et al. 2021; this study). A more recent discovery of nodulating symbionts in a species previously described as non-symbiotic is *P. azotifigens*, originally isolated from rice paddy soils in South Korea by Choi and Im (2018). This species is now revealed to be a common symbiont of widespread *Mimosa* species in Brazil and Kenya, but also of other neotropical mimosoids (*Jupunba*, *Parapiptadenia*). Intriguingly, although the four genome-sequenced symbiotic *P. azotifigens* strains (JPY530, JPY534, JPY105, and UYPR1.413) all harbor *nif* genes (Taulé et al. 2012; de Meyer et al. 2015b; this study), the genome of the non-symbiotic type strain, NF2-5-3^T^, despite its epithet, does not (data not shown).

As reported by the earlier study of Bontemps et al. (2010), and now with the addition of 90 new strains, there was relatively little evidence of HGT of symbiosis-related genes between the three main *Mimosa*-nodulating *Paraburkholderia* 16S rRNA-*recA* lineages. Indeed, the *nodC* and *nifH* sequences of most strains were congruent with their 16S rRNA-*recA* sequences, with the three largest SCs (SC2-3, SC4-5, and SC6) largely reconstructed in their phylogenies; these *nodC*- and *nifH*-derived lineages coincided with the “symbiovars” recently described by Paulitsch et al. (2020b) for *Mimosa*-nodulating *Paraburkholderia*. Nevertheless, there was some HGT of the symbiosis-determining loci observed between the SCs, which is to be expected as they often share the same habitats and even the same host plants, and all strains tested can nodulate at least one “common” host, *M. pudica*, regardless of which SC they belong to (Bontemps et al. 2010; this study). The possibility of inter-SC HGT had already been suggested by a study of invasive *Mimosa* symbionts in China in which *P. mimosarum* strains harbored *P. phymatum nodA* genes (Liu et al. 2012), and by a study of Brazilian *Calliandra* symbionts that were mainly *P. nodosa* in terms of their core genomes, but harbored *P. atlantica*/*P. youngii*-type *nodC* genes (Silva et al. 2018).

### 4.2. Cupriavidus is an important nodulating symbiont of widespread Mimosa species in Brazil

Although symbiotic *C. necator*-like strains were isolated from nodules of common bean and *Leucaena* plants used to trap rhizobia from pastures in MG in central Brazil by da Silva et al. (2012), the present study is the first report of Brazilian *Mimosa* species being nodulated in the field by *Cupriavidus*. This is surprising given that *Cupriavidus* has long been a major component of studies of *Mimosa* from elsewhere in its native and invasive range (Chen et al. 2001, 2003a, 2005b; Barrett and Parker 2005, 2006; Liu et al. 2012; Klonowska et al. 2012; Mishra et al. 2012; Gehlot et al. 2013; Reeve et al. 2015; Platero et al. 2016). However, most studies in Brazil (e.g. Bontemps et al. 2010) have focused primarily on the enormous variety of endemic *Mimosa* species in the highlands of central Brazil within the Cerrado and Caatinga biomes, as well as the *Campo rupestre* environments that arise from these biomes at altitudes above 900 m; the highly acidic and nutrient-poor soils of *Campo rupestre* environments are not only known drivers of endemism (Simon and Proença 2000), but will also favor acid-tolerant *Paraburkholderia* as potential symbionts (Stopnisek et al. 2014). Therefore, a probable reason for the previous absence of reports of *Cupriavidus* as a *Mimosa* symbiont was that endemic Brazilian *Mimosa* spp. generally do not encounter these bacteria in their natural habitats. They also tend to have a reduced ability to nodulate with *Cupriavidus* strains in *ex situ* single strain inoculation trials, suggesting that they are symbiotically incompatible partners (Elliott et al. 2007a; dos Reis Junior et al. 2010).

Most of the *Cupriavidus* strains in the present study were isolated from *M. pudica*, but also from *M. tenuiflora* and *M. candollei* (syn. *M. quadrivalvis*), growing in disturbed areas, pastures, and roadsides in the states of CE, ES, MG, RJ, RR. Genotypically, the strains were remarkably uniform, as almost all were *C. taiwanensis*, albeit with *C. necator*-like *nodC* and *nifH* genes like those of the symbionts of endemic Uruguayan *Mimosa* species (Platero et al. 2016; Iriarte et al. 2016). Such an unusual combination of core and symbiosis genes has not been previously observed in any native or invasive environment in which *Cupriavidus* has been isolated as a *Mimosa* symbiont (Chen et al. 2003a, 2005b; Andam et al. 2007; Mishra et al. 2012; Liu et al. 2012, 2020; Andrus et al. 2012; Klonowska et al. 2012; Gehlot et al. 2013; Parker, 2015). Compared to *Paraburkholderia* there are very few species of symbiotic *Cupriavidus*. Furthermore, the nodulating species have a remarkable monophyly of sym-genes, with most being close to either *C. taiwanensis* (Chen et al. 2003a, 2005b; Andam et al. 2007; Andrus et al. 2012; Klonowska et al. 2012; Mishra et al. 2012; Parker 2015) or to *C. necator* (da Silva et al. 2012; Taulé et al. 2012; Platero et al. 2016; this study). *Cupriavidus taiwanensis*, *C. neocaledonicus* and *C. necator* are the only validly named species confirmed as symbionts to date (Chen et al. 2001, 2003b; da Silva et al. 2012; Klonowska et al. 2020), although others are likely to be described. For example, *C. necator*- and *C. pinatubonensis*-like strains were the dominant symbionts of endemic *Mimosa* species from the heavy metal-rich mining area of Minas in Uruguay (Platero et al. 2016), and these probably represent new species. Meanwhile, the present study suggests a novel *Cupriavidus* species close to *C. lacunae* comprising UYPR2.512, a symbiont of *Parapiptadenia rigida* from Uruguay (Taulé et al. 2012; De Meyer et al. 2015a), and BR13378 from *M. pudica* in RJ (this study). Moreover, JPY540 isolated from *M. pudica* in Puerto Rico is closest to, but divergent from *C. oxalaticus*, and hence may also represent a novel species.

### 4.3. Factors driving Beta-rhizobial diversity in Brazil

Host endemism (altitude, location), and the highly location-specific edaphic factors that are associated with endemism (Simon and Proença 2000), have most likely driven co-evolution between endemic/biome-restricted *Mimosa* spp. and their symbionts. In the largest center of radiation of *Mimosa* in central Brazil, this is revealed as an almost exclusive association with *Paraburkholderia*, particularly *P. youngii* in SC6, and 16S rRNA-*recA* Groups K, L, M, and O in SC4-5 (Bontemps et al. 2010; this study), or occasionally with the much rarer but related genus, *Trinickia*, which has so far only been isolated from *M. cordistipula* and *M. misera* in BA (Bontemps et al. 2010; Estrada de Los Santos et al. 2018). It contrasts with the second largest *Mimosa* diversity center, Central Mexico, where endemic *Mimosa* spp. are mostly nodulated by *Rhizobium* or *Sinorhizobium* (Bontemps et al. 2016). A significant difference between central Brazil and central Mexico is that soils are highly acidic in the former (dos Reis Junior et al. 2010) and neutral-alkaline in the latter (Bontemps et al. 2016). Low soil pH is a known factor in promoting the occurrence of acid-tolerant *Paraburkholderia* (Stopnisek et al. 2014) as symbionts in both the main centers of Beta-rhizobial diversity, central Brazil and the CCR of South Africa (Garau et al. 2009; Howieson et al. 2013; Lemaire et al. 2015, 2016; Pires et al. 2018), especially when combined with low soil fertility (Elliott et al. 2009; Pires et al. 2018; Soares-Neto et al. 2022).

Therefore, the pattern that we now see in central Brazil is that native/endemic *Mimosa* spp. are largely nodulated by diverse genotypes of *Paraburkholderia* that they have co-evolved with in these acidic, nutrient-poor and high-Al soils (Simon and Proença 2000), whereas the widespread *Mimosa* spp. that are common in lowland Brazil (but also in disturbed areas in other parts of the country) are nodulated by *Paraburkholderia*, *Cupriavidus* and *Rhizobium* strains, depending on the sampling location. An RDA with *M. pudica*, the most widely sampled *Mimosa* host in the present study, revealed that various edaphic factors like soil pH, Ca, and P were important for separating the three bacterial genera. As stated above, low soil pH (and possibly high levels of potentially toxic aluminum ions, Al^3+^) was associated with a preference for *Paraburkholderia*. However, although alkaline pH has previously been shown to favor *Cupriavidus* (Mishra et al. 2012), in the present study this was not so obvious, perhaps because none of the sampled soils could be classified as such. Indeed, the soils containing *Cupriavidus* were mildly acidic with none above pH 6.8, suggesting that other edaphic factors are at least as crucial as elevated pH for explaining the presence of *Cupriavidus* symbionts, such as P, Mg, K, fertility, and the occurrence of heavy metals (Elliott et al. 2009; Mishra et al. 2012; Klonowska et al. 2012; Gehlot et al. 2013; Platero et al. 2016; Liu et al. 2012, 2020). That being said, it is clear that above-neutral pH is essential for the presence of *Mimosa*-nodulating *Rhizobium*, as evidenced by studies from Brazil (Baraúna et al. 2016; Pires et al. 2018; this study), Mexico (Bontemps et al. 2016), and China (Liu et al. 2020), amongst others.

## 5. Conclusion

There thus appear to be two stories associated with Brazilian *Mimosa* symbionts: that of the *Paraburkholderia* (and *Trinickia*) symbionts of upland endemics that grow in acidic, low nutrient and high Al-containing soils, and that of the *Paraburkholderia*, *Cupriavidus* and *Rhizobium* symbionts of lowland widespread species that grow in highly varied soils, and which select their symbionts according to those that are dominant in any particular soil according to its chemistry (pH, organic matter, fertility, nutrient status). It is also not surprising that the pantropical invasive *Mimosa* spp., such as *M. diplotricha*, *M. pigra* and *M. pudica*, are all typical widespread species in their native neotropics, and this might also explain why the same few “typical” symbionts of these species (*e.g.*, *C. taiwanensis*, *P. mimosarum* and *P. phymatum*) are isolated repeatedly in their invasive environments (Chen et al. 2003a, 2005; Andrus et al. 2012; Liu et al. 2012, 2020; Klonowska et al. 2012; Gehlot et al. 2013; Melkonian et al. 2014).

It is also now clear that Brazil and the broader neotropics and neo-subtropics constitute a significant habitat for a diverse range of Beta-rhizobia that are the natural nodulating symbionts of not only *Mimosa*, but also of other mimosoid genera, such as *Anadenanthera, Calliandra*, *Jupunba*, *Lachesiodendron*, *Parapiptadenia*, *Piptadenia*, and *Pseudopiptadenia*, and that similar genotypes of *Paraburkholderia* and *Cupriavidus* that nodulate *Mimosa* spp. also nodulate these hosts (Taulé et al. 2012; Bournaud et al. 2013, Silva et al. 2018; Zilli et al. 2021). The recent discovery that the non-mimosoid Caesalpinioideae species, *Chamaecrista eitenorum*, is nodulated effectively by *P. nodosa*-like strains (Casaes et al. 2024) further indicates that it is likely that we are only scratching the surface of the actual diversity of neotropical and sub-neotropical Beta-rhizobial-legume interactions. Indeed, given the propensity of South American *Paraburkholderia* and *Cupriavidus* strains to nodulate effectively with papilionoids like common bean and Siratro (*Macroptilium atropurpureum*) (da Silva et al. 2012; Dall-Agnol et al. 2016, 2017) it is highly probable that Beta-rhizobia will be isolated from more crop legumes, as well as from their diverse native neotropical relatives.

## Data availability

All data are available in this manuscript and in the supplementary materials.

## Acknowledgements

We thank CNPq (Brazilian National Council for Scientific and Technological Development) for productivity grants awarded to some authors; CAPES (Coordination of Superior Level Staff Improvement) for student grants and for a grant to EKJ via the Ciência sem Fronteiras program, and to FAPERJ for the fellowship “Cientista do Nosso Estado”. The James Hutton Institute is supported by the Rural & Environment Science & Analytical Services (RESAS), a division of the Scottish Government. Unless noted otherwise, genome sequencing was provided by MicrobesNG (http://www.microbesng.uk), which is supported by the BBSRC (grant number BB/L024209/1).

---

**Appendix A. Supplementary data**

**Fig. S1.**
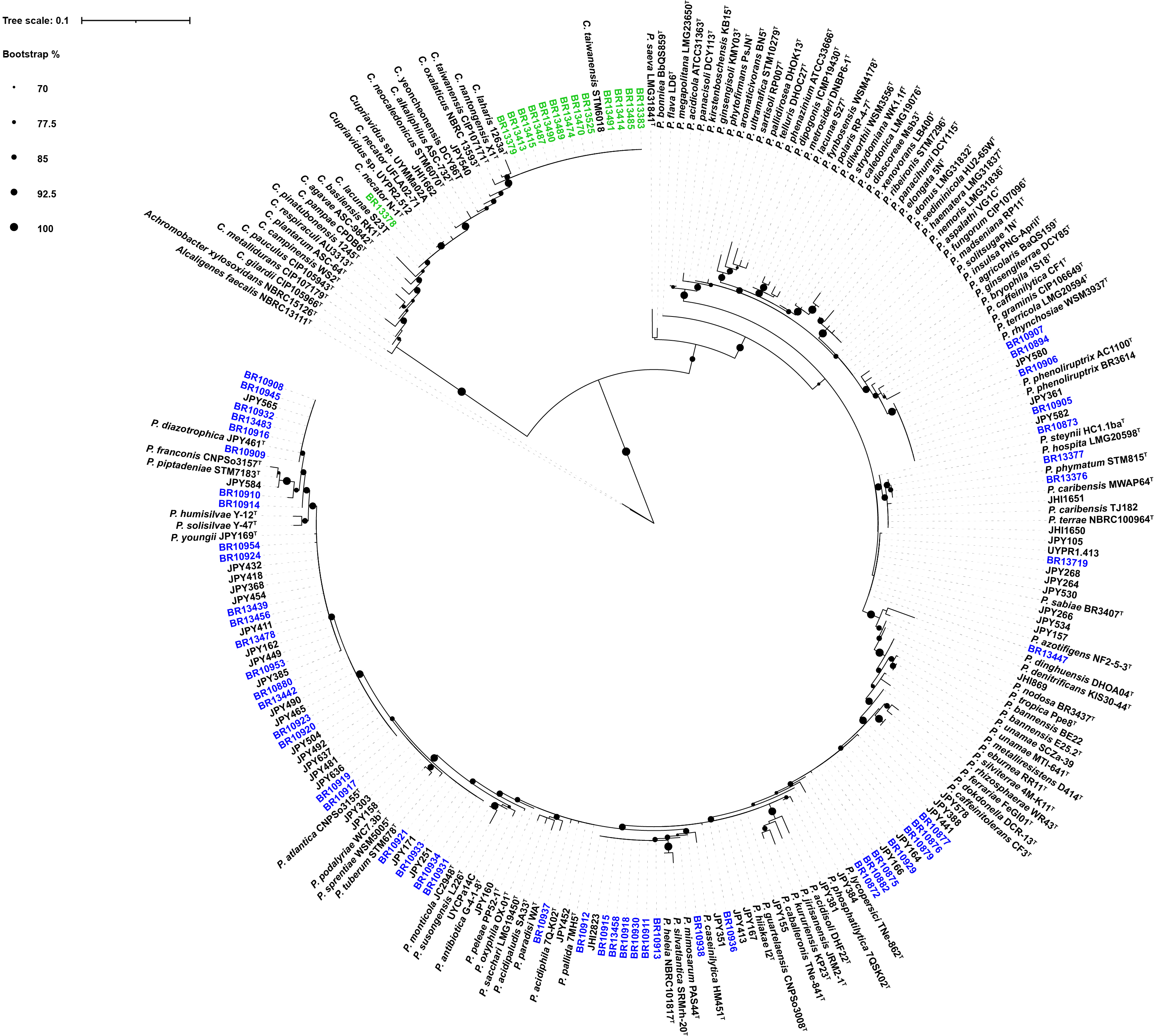
16S rRNA phylogeny of *Paraburkholderia* and *Cupriavidus*. **Maximum-likelihood** tree built using IQ-TREE, with ultrafast bootstrap analysis (1000 iterations) and the ‘TIM2+F+I+G4’ best-fit model according to Bayesian Information Criterion. The alignment contained 243 sequences and 850 nucleotide positions with 120 parsimony informative sites.

**Fig. S2.**
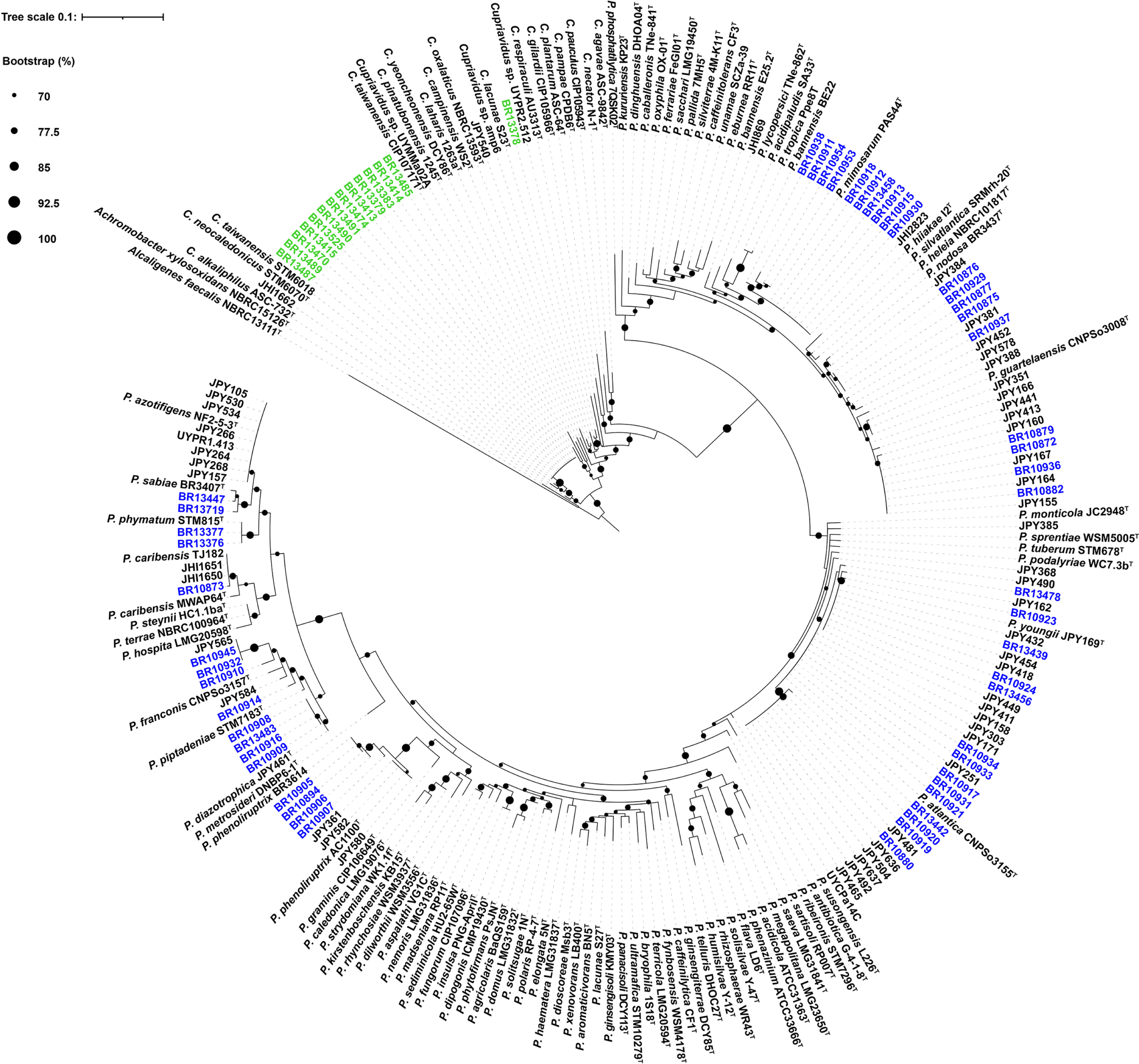
*recA* phylogeny of *Paraburkholderia* and *Cupriavidus*. **Maximum-likelihood** tree built using IQ-TREE, with ultrafast bootstrap analysis (1000 iterations) and the ‘TIM2+F+I+G4’ best-fit model according to Bayesian Information Criterion. The alignment contained 226 sequences and 339 nucleotide positions with 131 parsimony informative sites.

**Fig. S3.**
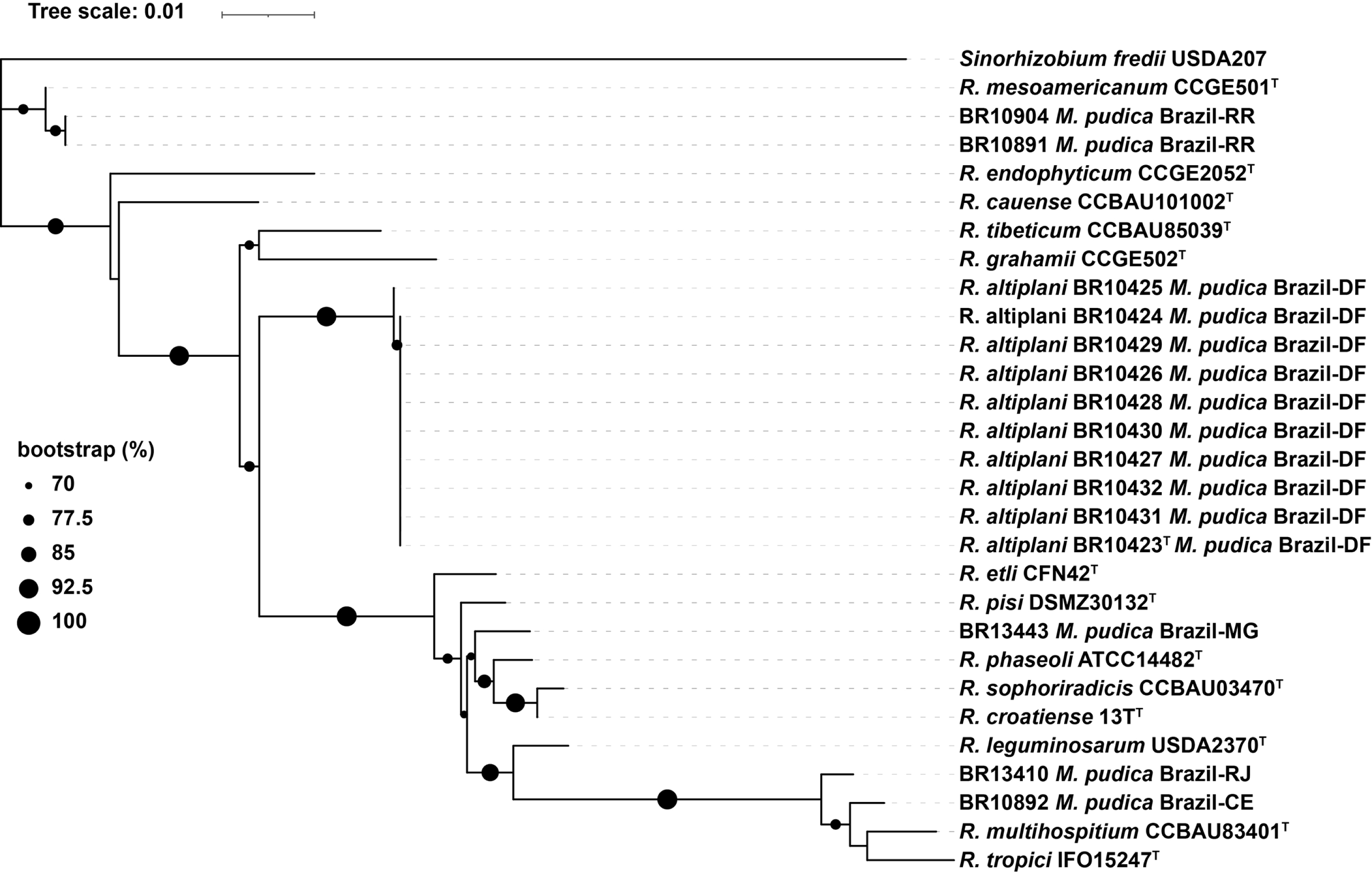
16S rRNA phylogeny of *Rhizobium*. **Maximum-likelihood** tree built using IQ-TREE, with ultrafast bootstrap analysis (1000 iterations) and the ‘TN+F+I+G4’ best-fit model according to Bayesian Information Criterion. The alignment contained 29 sequences and 1484 nucleotide positions with 101 parsimony-informative sites.

**Table S1** Rhizobial and reference strains examined in this study, their geographical origins, host plants, and gene and genome sequence accession numbers.

**Table S2** ANIb analysis of genomes sequenced from selected *Paraburkholderia* (a) and *Cupriavidus* (b) strains compared to the nearest type strains and to other reference strains.

## Data availability

All data supporting the findings of this study are available within the paper and within its supplementary data published online.

## References

Achouak, W., 1999. *Burkholderia caribensis* sp. nov., an exopolysaccharide-producing bacterium isolated from vertisol microaggregates in Martinique. International Journal of Systematic Bacteriology 49, 787–794. doi:10.1099/00207713-49-2-787

Andam, C.P., Mondo, S.J., Parker, M.A., 2007. Monophyly of *nodA* and *nifH* genes across Texan and Costa Rican populations of *Cupriavidus* nodule symbionts. Applied and Environmental Microbiology 73, 4686–4690. doi:10.1128/AEM.00160-07

Andrews, M., Andrews, M.E., 2017. Specificity in legume-rhizobia symbioses. International Journal of Molecular Sciences 18. doi:10.3390/IJMS18040705

Andrus, A.D., Andam, C., Parker, M.A., 2012. American origin of *Cupriavidus* bacteria associated with invasive *Mimosa* legumes in the Philippines. FEMS Microbiology Ecology 80, 747–750. doi:10.1111/J.1574-6941.2012.01342.X

Arahal, D.R., 2014. Whole-genome analyses: Average nucleotide identity. Methods in Microbiology 41, 103–122. doi:10.1016/BS.MIM.2014.07.002

Ardley, J., Sprent, J., 2021. Evolution and biogeography of actinorhizal plants and legumes: A comparison. Journal of Ecology 109, 1098–1121. doi:10.1111/1365-2745.13600

Bankevich, A., Nurk, S., Antipov, D., Gurevich, A.A., Dvorkin, M., Kulikov, A.S., Lesin, V.M., Nikolenko, S.I., Pham, S., Prjibelski, A.D., Pyshkin, A.V., Sirotkin, A.V., Vyahhi, N., Tesler, G., Alekseyev, M.A., Pevzner, P.A., 2012. SPAdes: A new genome assembly algorithm and its applications to single-cell sequencing. Journal of Computational Biology 19, 455–477. doi:10.1089/CMB.2012.0021

Baraúna, A.C., Rouws, L.F.M., Simoes-Araujo, J.L., dos Reis Junior, F.B., Iannetta, P.P.M., Maluk, M., Goi, S.R., Reis, V.M., James, E.K., Zilli, J.E., 2016. *Rhizobium altiplani* sp. nov., isolated from effective nodules on *Mimosa pudica* growing in untypically alkaline soil in central Brazil. International Journal of Systematic and Evolutionary Microbiology 66, 4118–4124. doi:10.1099/IJSEM.0.001322

Barrett, C.F., Parker, M.A., 2005. Prevalence of *Burkholderia* sp. nodule symbionts on four mimosoid legumes from Barro Colorado Island, Panama. Systematic and Applied Microbiology 28, 57–65. doi:10.1016/J.SYAPM.2004.09.002

Benson, D.A., Cavanaugh, M., Clark, K., Karsch-Mizrachi, I., Lipman, D.J., Ostell, J., Sayers, E.W., 2017. GenBank. Nucleic Acids Research 45, D37–D42. doi:10.1093/NAR/GKW1070

Beukes, C.W., Palmer, M., Manyaka, P., Chan, W.Y., Avontuur, J.R., van Zyl, E., Huntemann, M., Clum, A., Pillay, M., Palaniappan, K., Varghese, N., Mikhailova, N., Stamatis, D., Reddy, T.B.K., Daum, C., Shapiro, N., Markowitz, V., Ivanova, N., Kyrpides, N., Woyke, T., Blom, J., Whitman, W.B., Venter, S.N., Steenkamp, E.T., 2017. Genome data provides high support for generic boundaries in *Burkholderia* sensu lato. Frontiers in Microbiology 8. doi:10.3389/FMICB.2017.01154

Beukes, C.W., Venter, S.N., Law, I.J., Phalane, F.L., Steenkamp, E.T., 2013. South African papilionoid legumes are nodulated by diverse *Burkholderia* with unique nodulation and nitrogen-fixation loci. PLoS ONE 8. doi:10.1371/JOURNAL.PONE.0068406

Bontemps, C., Elliott, G.N., Simon, M.F., dos Reis Junior, F.B., Gross, E., Lawton, R.C., Neto, N.E., De FÁtima Loureiro, M., De Faria, S.M., Sprent, J.I., James, E.K., Young, J.P.W., 2010. Burkholderia species are ancient symbionts of legumes. Molecular Ecology 19, 44–52. doi:10.1111/J.1365-294X.2009.04458.X

Bontemps, C., Rogel, M.A., Wiechmann, A., Mussabekova, A., Moody, S., Simon, M.F., Moulin, L., Elliott, G.N., Lacercat-Didier, L., Dasilva, C., Grether, R., Camargo-Ricalde, S.L., Chen, W., Sprent, J.I., Martínez-Romero, E., Young, J.P.W., James, E.K., 2016. Endemic Mimosa species from Mexico prefer alphaproteobacterial rhizobial symbionts. New Phytologist 209, 319–333. doi:10.1111/NPH.13573

Bournaud, C., de Faria, S.M., dos Santos, J.M.F., Tisseyre, P., Silva, M., Chaintreuil, C., Gross, E., James, E.K., Prin, Y., Moulin, L., 2013. *Burkholderia* species are the most common and preferred nodulating symbionts of the *Piptadenia* group (Tribe Mimoseae). PLoS ONE 8. doi:10.1371/JOURNAL.PONE.0063478

Bournaud, C., Moulin, L., Cnockaert, M., de Faria, S., Prin, Y., Severac, D., Vandamme, P., 2017. *Paraburkholderia piptadeniae* sp. nov. and *Paraburkholderia ribeironis* sp. nov., two root-nodulating symbiotic species of *Piptadenia gonoacantha* in Brazil. International Journal of Systematic and Evolutionary Microbiology 67, 432–440. doi:10.1099/IJSEM.0.001648

dos Reis Junior, F.B., Simon, M.F., Gross, E., Boddey, R.M., Elliott, G.N., Neto, N.E., de Fatima Loureiro, M., de Queiroz, L.P., Scotti, M.R., Chen, W.M., Norén, A., Rubio, M.C., de Faria, S.M., Bontemps, C., Goi, S.R., Young, J.P.W., Sprent, J.I., James, E.K., 2010. Nodulation and nitrogen fixation by *Mimosa* spp. in the Cerrado and Caatinga biomes of Brazil. New Phytologist 186, 934–946. doi:10.1111/J.1469-8137.2010.03267.X

Casaes, P.A., Ferreira dos Santos, J.M., Silva, V.C., Rhem, M.F.K., Teixeira Cota, M.M., de Faria, S.M., Rando, J.G., James, E.K., Gross, E., 2024. The radiation of nodulated *Chamaecrista* species from the rainforest into more diverse habitats has been accompanied by a reduction in growth form and a shift from fixation threads to symbiosomes. Journal of Experimental Botany. doi:10.1093/JXB/ERAE134

Chen, W.M., de Faria, S.M., James, E.K., Elliott, G.N., Lin, K.Y., Chou, J.H., Sheu, S.Y., Cnockaert, M., Sprent, J.I., Vandamme, P., 2007. *Burkholderia nodosa* sp. nov., isolated from root nodules of the woody Brazilian legumes *Mimosa bimucronata* and *Mimosa scabrella*. International Journal of Systematic and Evolutionary Microbiology 57, 1055–1059. doi:10.1099/IJS.0.64873-0

Chen, W.M., De Faria, S.M., Straliotto, R., Pitard, R.M., Simões-Araùjo, J.L., Chou, J.H., Chou, Y.J., Barrios, E., Prescott, A.R., Elliott, G.N., Sprent, J.I., Young, J.P.W., James, E.K., 2005a. Proof that *Burkholderia* strains form effective symbioses with legumes: A study of novel *Mimosa*-nodulating strains from South America. Applied and Environmental Microbiology 71, 7461–7471. doi:10.1128/AEM.71.11.7461-7471.2005

Chen, W.M., de Farja, S.M., Chou, J.H., James, E.K., Elliott, G.N., Sprent, J.I., Bontemps, C., Young, J.P.W., Vandamme, P., 2008. *Burkholderia sabiae* sp. nov., isolated from root nodules of *Mimosa caesalpiniifolia*. International Journal of Systematic and Evolutionary Microbiology 58, 2174–2179. doi:10.1099/IJS.0.65816-0

Chen, W.M., James, E.K., Chou, J.H., Sheu, S.Y., Yang, S.Z., Sprent, J.I., 2005b. β-rhizobia from *Mimosa pigra*, a newly discovered invasive plant in Taiwan. New Phytologist 168, 661–675. doi:10.1111/J.1469-8137.2005.01533.X

Chen, W.M., James, E.K., Coenye, T., Chou, J.H., Barrios, E., de Faria, S.M., Elliott, G.N., Sheu, S.Y., Sprent, J.I., Vandamme, P., 2006. *Burkholderia mimosarum* sp. nov., isolated from root nodules of Mimosa spp. from Taiwan and South America. International Journal of Systematic and Evolutionary Microbiology 56, 1847–1851. doi:10.1099/IJS.0.64325-0

Chen, W.M., James, E.K., Prescott, A.R., Kierans, M., Sprent, J.I., 2003a. Nodulation of *Mimosa* spp. by the β-Proteobacterium *Ralstonia taiwanensis*. Molecular Plant-Microbe Interactions 16, 1051–1061. doi:10.1094/MPMI.2003.16.12.1051

Chen, W.M., Laevens, S., Lee, T.M., Coenye, T., De Vos, P., Mergeay, M., Vandamme, P., 2001. *Ralstonia taiwanensis* sp. nov., isolated from root nodules of *Mimosa* species and sputum of a cystic fibrosis patient. International Journal of Systematic and Evolutionary Microbiology 51, 1729–1735. doi:10.1099/00207713-51-5-1729

Chen, W.M., Moulin, L., Bontemps, C., Vandamme, P., Béna, G., Boivin-Masson, C., 2003b. Legume symbiotic nitrogen fixation by β-proteobacteria is widespread in nature. Journal of Bacteriology 185, 7266–7272. doi:10.1128/JB.185.24.7266-7272.2003

Choi, G.M., Im, W.T., 2018. *Paraburkholderia azotifigens* sp. nov., a nitrogen-fixing bacterium isolated from paddy soil. International Journal of Systematic and Evolutionary Microbiology 68, 310–316. doi:10.1099/IJSEM.0.002505

Coenye, T., Henry, D., Speert, D.P., Vandamme, P., 2004. *Burkholderia phenoliruptrix* sp. nov., to accommodate the 2,4,5-trichlorophenoxyacetic acid and halophenol-degrading strain AC1100. Systematic and Applied Microbiology 27, 623–627. doi:10.1078/0723202042369992

Cunha, C. de O., Zuleta, L.F.G., de Almeida, L.G.P., Ciapina, L.P., Borges, W.L., Pitard, R.M., Baldani, J.I., Straliotto, R., de Faria, S.M., Hungria, M., Cavada, B.S., Mercante, F.M., de Vasconcelos, A.T.R., 2012. Complete genome sequence of *Burkholderia phenoliruptrix* BR3459a (CLA1), a heat-tolerant, nitrogen-fixing symbiont of *Mimosa flocculosa*. Journal of Bacteriology 194, 6675–6676. doi:10.1128/JB.01821-12

da Silva, K., Florentino, L.A., da Silva, K.B., de Brandt, E., Vandamme, P., de Souza Moreira, F.M., 2012. *Cupriavidus necator* isolates are able to fix nitrogen in symbiosis with different legume species. Systematic and Applied Microbiology 35, 175–182. doi:10.1016/J.SYAPM.2011.10.005

Dall’Agnol, R.F., Bournaud, C., de Faria, S.M., Béna, G., Moulin, L., Hungria, M., 2017. Genetic diversity of symbiotic *Paraburkholderia* species isolated from nodules of *Mimosa pudica* (L.) and *Phaseolus vulgaris* (L.) grown in soils of the Brazilian Atlantic Forest (Mata Atlântica). FEMS Microbiology Ecology 93. doi:10.1093/FEMSEC/FIX027

Dall’Agnol, R.F., Plotegher, F., Souza, R.C., Mendes, I.C., Dos Reis, F.B., Béna, G., Moulin, L., Hungria, M., 2016. *Paraburkholderia nodosa* is the main N_2_-fixing species trapped by promiscuous common bean (*Phaseolus vulgaris* L.) in the Brazilian “Cerradão.” FEMS Microbiology Ecology 92. doi:10.1093/FEMSEC/FIW108

de Castro Pires, R., dos Reis Junior, F.B., Zilli, J.E., Fischer, D., Hofmann, A., James, E.K., Simon, M.F., 2018. Soil characteristics determine the rhizobia in association with different species of *Mimosa* in central Brazil. Plant and Soil 423, 411–428. doi:10.1007/S11104-017-3521-5

de Faria, S.M., Ringelberg, J.J., Gross, E., Koenen, E.J.M., Cardoso, D., Ametsitsi, G.K.D., Akomatey, J., Maluk, M., Tak, N., Gehlot, H.S., Wright, K.M., Teaumroong, N., Songwattana, P., de Lima, H.C., Prin, Y., Zartman, C.E., Sprent, J.I., Ardley, J., Hughes, C.E., James, E.K., 2022. The innovation of the symbiosome has enhanced the evolutionary stability of nitrogen fixation in legumes. New Phytologist 235, 2365–2377. doi:10.1111/NPH.18321

De Meyer, S.E., Fabiano, E., Tian, R., Van Berkum, P., Seshadri, R., Reddy, T.B.K., Markowitz, V., Ivanova, N., Pati, A., Woyke, T., Howieson, J., Kyrpides, N., Reeve, W., 2015a. High-quality permanent draft genome sequence of the *Parapiptadenia rigida*-nodulating *Burkholderia* sp. strain UYPR1.413. Standards in Genomic Sciences 10. doi:10.1186/S40793-015-0018-9

De Meyer, S.E., Fabiano, E., Tian, R., Van Berkum, P., Seshadri, R., Reddy, T.B.K., Markowitz, V., Ivanova, N.N., Pati, A., Woyke, T., Howieson, J., Kyrpides, N.C., Reeve, W., 2015b. High-quality permanent draft genome sequence of the *Parapiptadenia rigida*-nodulating *Cupriavidus* sp. strain UYPR2.512. Standards in Genomic Sciences 10. doi:10.1186/1944-3277-10-13

De Meyer, S.E., Parker, M., Van Berkum, P., Tian, R., Seshadri, R., Reddy, T.B.K., Markowitz, V., Ivanova, N., Pati, A., Woyke, T., Kyrpides, N., Howieson, J., Reeve, W., 2015c. High-quality permanent draft genome sequence of the *Mimosa asperata* - nodulating *Cupriavidus* sp. strain AMP6. Standards in Genomic Sciences 10. doi:10.1186/S40793-015-0074-1

Dias, M.A.M., Bomfim, C.S.G., Rodrigues, D.R., da Silva, A.F., Santos, J.C.S., do Nascimento, T.R., Martins, L.M.V., Dantas, B.F., Ribeiro, P.R. de A., de Freitas, A.D.S., Fernandes-Júnior, P.I., 2021. *Paraburkholderia* spp. are the main rhizobial microsymbionts of *Mimosa tenuiflora* (Willd.) Poir. in soils of the Brazilian tropical dry forests (Caatinga biome). Systematic and Applied Microbiology 44. doi:10.1016/J.SYAPM.2021.126208

Dludlu, Meshack Nkosinathi, Chimphango, S.B.M., Stirton, C.H., Muasya, A.M., 2018. Differential preference of *Burkholderia* and *Mesorhizobium* to pH and soil types in the Core Cape Subregion, South Africa. Genes 9. doi:10.3390/GENES9010002

Dludlu, M. N., Chimphango, S.B.M., Walker, G., Stirton, C.H., Muasya, A.M., 2018. Horizontal gene transfer among rhizobia of the Core Cape Subregion of southern Africa. South African Journal of Botany 118, 342–352. doi:10.1016/J.SAJB.2018.02.406

Dutra, V.F., Morales, M., Jordão, L.S.B., Borges, L.M., Silveira, F.S., Simon, M.F., Santos-Silva, J., Nascimento, J.G.A., Ribas, O.D.S., 2020. *Mimosa* in Flora do Brasil 2020. Jardim Botânico do Rio de Janeiro. https://floradobrasil2020.jbrj.gov.br/FB23084

Edgar, R.C., 2004. MUSCLE: A multiple sequence alignment method with reduced time and space complexity. BMC Bioinformatics 5. doi:10.1186/1471-2105-5-113

Elliott, G.N., Chen, W.M., Bontemps, C., Chou, J.H., Young, J.P.W., Sprent, J.I., James, E.K., 2007a. Nodulation of *Cyclopia* spp. (Leguminosae, Papilionoideae) by Burkholderia tuberum. Annals of Botany 100, 1403–1411. doi:10.1093/AOB/MCM227

Elliott, G.N., Chen, W.M., Chou, J.H., Wang, H.C., Sheu, S.Y., Perin, L., Reis, V.M., Moulin, L., Simon, M.F., Bontemps, C., Sutherland, J.M., Bessi, R., De Faria, S.M., Trinick, M.J., Prescott, A.R., Sprent, J.I., James, E.K., 2007b. *Burkholderia phymatum* is a highly effective nitrogen-fixing symbiont of *Mimosa* spp. and fixes nitrogen ex planta. New Phytologist 173, 168–180. doi:10.1111/J.1469-8137.2006.01894.X

Elliott, G.N., Chou, J.H., Chen, W.M., Bloemberg, G.V., Bontemps, C., Martínez-Romero, E., Velázquez, E., Young, J.P.W., Sprent, J.I., James, E.K., 2009. *Burkholderia* spp. are the most competitive symbionts of *Mimosa*, particularly under N-limited conditions. Environmental Microbiology 11, 762–778. doi:10.1111/J.1462-2920.2008.01799.X

Estrada-de los Santos, P., Palmer, M., Chávez-Ramírez, B., Beukes, C., Steenkamp, E.T., Briscoe, L., Khan, N., Maluk, M., Lafos, M., Humm, E., Arrabit, M., Crook, M., Gross, E., Simon, M.F., dos Reis Junior, F.B., Whitman, W.B., Shapiro, N., Poole, P.S., Hirsch, A.M., Venter, S.N., James, E.K., 2018. Whole genome analyses suggests that *Burkholderia* sensu lato contains two additional novel genera (*Mycetohabitans* gen. nov., and *Trinickia* gen. nov.): Implications for the evolution of diazotrophy and nodulation in the Burkholderiaceae. Genes 9. doi:10.3390/GENES9080389

Fred, E.B., Waksman, S.A., 1928. Laboratory manual of general microbiology: with special reference to the microorganisms of the soil. New York, McGraw-Hill book Company, Incorporated.

Garau, G., Yates, R.J., Deiana, P., Howieson, J.G., 2009. Novel strains of nodulating *Burkholderia* have a role in nitrogen fixation with papilionoid herbaceous legumes adapted to acid, infertile soils. Soil Biology and Biochemistry 41, 125–134. doi:10.1016/J.SOILBIO.2008.10.011

Gehlot, H.S., Tak, N., Kaushik, M., Mitra, S., Chen, W.M., Poweleit, N., Panwar, D., Poonar, N., Parihar, R., Tak, A., Sankhla, I.S., Ojha, A., Rao, S.R., Simon, M.F., Reis Junior, F.B.D., Perigolo, N., Tripathi, A.K., Sprent, J.I., Young, J.P.W., James, E.K., Gyaneshwar, P., 2013. An invasive *Mimosa* in India does not adopt the symbionts of its native relatives. Annals of Botany 112, 179–196. doi:10.1093/AOB/MCT112

Goris, J., Konstantinidis, K.T., Klappenbach, J.A., Coenye, T., Vandamme, P., Tiedje, J.M., 2007. DNA-DNA hybridization values and their relationship to whole-genome sequence similarities. International Journal of Systematic and Evolutionary Microbiology 57, 81–91. doi:10.1099/IJS.0.64483-0

Gyaneshwar, P., Hirsch, A.M., Moulin, L., Chen, W.M., Elliott, G.N., Bontemps, C., De Los Santos, P.E., Gross, E., Dos Reis, F.B., Janet, I.S., Young, J.P.W., James, E.K., 2011. Legume-nodulating betaproteobacteria: Diversity, host range, and future prospects. Molecular Plant-Microbe Interactions 24, 1276–1288. doi:10.1094/MPMI-06-11-0172

Howieson, J.G., De Meyer, S.E., Vivas-Marfisi, A., Ratnayake, S., Ardley, J.K., Yates, R.J., 2013. Novel *Burkholderia* bacteria isolated from *Lebeckia ambigua* - A perennial suffrutescent legume of the fynbos. Soil Biology and Biochemistry 60, 55–64. doi:10.1016/J.SOILBIO.2013.01.009

Iriarte, A., Platero, R., Romero, V., Fabiano, E., Sotelo-Silveira, J.R., 2016. Draft genome sequence of *Cupriavidus* UYMMa02A, a novel beta-rhizobium species. Genome Announcements 4. doi:10.1128/GENOMEA.01258-16

Kalyaanamoorthy, S., Minh, B.Q., Wong, T.K.F., Von Haeseler, A., Jermiin, L.S., 2017. ModelFinder: Fast model selection for accurate phylogenetic estimates. Nature Methods 14, 587–589. doi:10.1038/NMETH.4285

Klepa, M.S., Janoni, V., Paulitsch, F., da Silva, A.R., do Carmo, M.R.B., Delamuta, J.R.M., Hungria, M., da Silva Batista, J.S., 2021. Molecular diversity of rhizobia-nodulating native *Mimosa* of Brazilian protected areas. Archives of Microbiology 203, 5533–5545. doi:10.1007/S00203-021-02537-7

Klonowska, A., Chaintreuil, C., Tisseyre, P., Miché, L., Melkonian, R., Ducousso, M., Laguerre, G., Brunel, B., Moulin, L., 2012. Biodiversity of *Mimosa* pudica rhizobial symbionts (*Cupriavidus taiwanensis*, Rhizobium mesoamericanum) in New Caledonia and their adaptation to heavy metal-rich soils. FEMS Microbiology Ecology 81, 618–635. doi:10.1111/J.1574-6941.2012.01393.X

Klonowska, A., Moulin, L., Ardley, J.K., Braun, F., Gollagher, M.M., Zandberg, J.D., Marinova, D.V., Huntemann, M., Reddy, T.B.K., Varghese, N.J., Woyke, T., Ivanova, N., Seshadri, R., Kyrpides, N., Reeve, W.G., 2020. Novel heavy metal resistance gene clusters are present in the genome of *Cupriavidus neocaledonicus* STM 6070, a new species of *Mimosa pudica* microsymbiont isolated from heavy-metal-rich mining site soil. BMC Genomics 21. doi:10.1186/S12864-020-6623-Z

Koeuth, T., Versalovic, J., Lupski, J.R., 1995. Differential subsequence conservation of interspersed repetitive *Streptococcus pneumoniae* BOX elements in diverse bacteria. Genome Research 5, 408–418. doi:10.1101/GR.5.4.408

Kumar, S., Stecher, G., Tamura, K., 2016. MEGA7: Molecular evolutionary genetics analysis version 7.0 for bigger datasets. Molecular Biology and Evolution 33, 1870–1874. doi:10.1093/MOLBEV/MSW054

Lammel, D.R., Cruz, L.M., Mescolotti, D., Stürmer, S.L., Cardoso, E.J.B.N., 2015. Woody Mimosa species are nodulated by *Burkholderia* in ombrophylous forest soils and their symbioses are enhanced by arbuscular mycorrhizal fungi (AMF). Plant and Soil 393, 123–135. doi:10.1007/S11104-015-2470-0

Langleib, M., Beracochea, M., Zabaleta, M., Battistoni, F., Sotelo-Silveira, J., Fabiano, E., Iriarte, A., Platero, R., 2019. Draft genome sequence of *Paraburkholderia* sp. UYCP14C, a rhizobium strain isolated from root nodules of *Calliandra parvifolia*. Microbiology Resource Announcements 8. doi:10.1128/MRA.00173-19

Lemaire, B., Chimphango, S.B.M., Stirton, C., Rafudeen, S., Honnay, O., Smets, E., Chen, W.M., Sprent, J., James, E.K., Muasya, A.M., 2016. Biogeographical patterns of legume-nodulating *Burkholderia* spp.: From African fynbos to continental scales. Applied and Environmental Microbiology 82, 5099–5115. doi:10.1128/AEM.00591-16

Lemaire, B., Dlodlo, O., Chimphango, S., Stirton, C., Schrire, B., Boatwright, J.S., Honnay, O., Smets, E., Sprent, J., James, E.K., Muasya, A.M., 2015. Symbiotic diversity, specificity and distribution of rhizobia in native legumes of the Core Cape Subregion (South Africa). FEMS Microbiology Ecology 91, 1–17. doi:10.1093/FEMSEC/FIU024

Letunic, I., Bork, P., 2021. Interactive tree of life (iTOL) v5: An online tool for phylogenetic tree display and annotation. Nucleic Acids Research 49, W293– W296. doi:10.1093/NAR/GKAB301

Liu, W.Y.Y., Ridgway, H.J., James, T.K., James, E.K., Chen, W.M., Sprent, J.I., Young, J.P.W., Andrews, M., 2014. *Burkholderia* sp. induces functional nodules on the South African invasive legume *Dipogon lignosus* (Phaseoleae) in New Zealand soils. Microbial Ecology 68, 542–555. doi:10.1007/S00248-014-0427-0

Liu, X., Wei, S., Wang, F., James, E.K., Guo, X., Zagar, C., Xia, L.G., Dong, X., Wang, Y.P., 2012. *Burkholderia* and *Cupriavidus* spp. are the preferred symbionts of *Mimosa* spp. in southern China. FEMS Microbiology Ecology 80, 417–426. doi:10.1111/J.1574-6941.2012.01310.X

Liu, X., You, S., Liu, H., Yuan, B., Wang, H., James, E.K., Wang, F., Cao, W., Liu, Z.K., 2020. Diversity and geographic distribution of microsymbionts associated with invasive *Mimosa* species in southern China. Frontiers in Microbiology 11. doi:10.3389/FMICB.2020.563389

L.P.W.G. The World Checklist of Vascular Plants (WCVP): Fabaceae, version 2023. V4 (R. Govaerts, Ed.). The Royal Botanic Gardens, Kew. 10.15468/mvhaj3.

Mavima, L., Beukes, C.W., Palmer, M., De Meyer, S.E., James, E.K., Maluk, M., Gross, E., dos Reis Junior, F.B., Avontuur, J.R., Chan, W.Y., Venter, S.N., Steenkamp, E.T., 2021. *Paraburkholderia youngii* sp. nov. and ‘*Paraburkholderia atlantica*’ – Brazilian and Mexican Mimosa-associated rhizobia that were previously known as *Paraburkholderia tuberum* sv. *mimosae*. Systematic and Applied Microbiology 44. doi:10.1016/J.SYAPM.2020.126152

Mavima, L., Beukes, C.W., Palmer, M., De Meyer, S.E., James, E.K., Maluk, M., Muasya, M.A., Avontuur, J.R., Chan, W.Y., Venter, S.N., Steenkamp, E.T., 2022. Delineation of *Paraburkholderia tuberum* sensu stricto and description of *Paraburkholderia podalyriae* sp. nov. nodulating the South African legume *Podalyria calyptrata*. Systematic and Applied Microbiology 45. doi:10.1016/J.SYAPM.2022.126316

Melkonian, R., Moulin, L., Béna, G., Tisseyre, P., Chaintreuil, C., Heulin, K., Rezkallah, N., Klonowska, A., Gonzalez, S., Simon, M., Chen, W.M., James, E.K., Laguerre, G., 2014. The geographical patterns of symbiont diversity in the invasive legume *Mimosa pudica* can be explained by the competitiveness of its symbionts and by the host genotype. Environmental Microbiology 16, 2099–2111. doi:10.1111/1462-2920.12286

Minh, B.Q., Nguyen, M.A.T., Von Haeseler, A., 2013. Ultrafast approximation for phylogenetic bootstrap. Molecular Biology and Evolution 30, 1188–1195. doi:10.1093/MOLBEV/MST024

Mishra, R.P.N., Tisseyre, P., Melkonian, R., Chaintreuil, C., Miché, L., Klonowska, A., Gonzalez, S., Bena, G., Laguerre, G., Moulin, L., 2012. Genetic diversity of *Mimosa pudica* rhizobial symbionts in soils of French Guiana: Investigating the origin and diversity of *Burkholderia phymatum* and other beta-rhizobia. FEMS Microbiology Ecology 79, 487–503. doi:10.1111/J.1574-6941.2011.01235.X

Norris, D.O., ‘t Mannetje, L., 1964. The symbiotic specialization of African *Trifolium* spp. in relation to their taxonomy and their agronomic use. East African Agricultural and Forestry Journal 29, 214–235. doi:10.1080/00128325.1964.11661928

Nurk, S., Bankevich, A., Antipov, D., Gurevich, A., Korobeynikov, A., Lapidus, A., Prjibelsky, A., Pyshkin, A., Sirotkin, A., Sirotkin, Y., Stepanauskas, R., McLean, J., Lasken, R., Clingenpeel, S.R., Woyke, T., Tesler, G., Alekseyev, M.A., Pevzner, P.A., 2013. Assembling genomes and mini-metagenomes from highly chimeric reads. Lecture Notes in Computer Science (Including Subseries Lecture Notes in Artificial Intelligence and Lecture Notes in Bioinformatics) 7821 LNBI, 158–170. doi:10.1007/978-3-642-37195-0_13

Parker, M.A., 2015. A single sym plasmid type predominates across diverse chromosomal lineages of *Cupriavidus* nodule symbionts. Systematic and Applied Microbiology 38, 417–423. doi:10.1016/J.SYAPM.2015.06.003

Parker, M.A., Wurtz, A.K., Paynter, Q., 2007. Nodule symbiosis of invasive *Mimosa* pigra in Australia and in ancestral habitats: A comparative analysis. Biological Invasions 9, 127–138. doi:10.1007/S10530-006-0009-2

Parte, A.C., Carbasse, J.S., Meier-Kolthoff, J.P., Reimer, L.C., Göker, M., 2020. List of prokaryotic names with standing in nomenclature (LPSN) moves to the DSMZ. International Journal of Systematic and Evolutionary Microbiology 70, 5607–5612. doi:10.1099/IJSEM.0.004332

Paulitsch, F., Dall’Agnol, R.F., Delamuta, J.R.M., Ribeiro, R.A., da Silva Batista, J.S., Hungria, M., 2020a. *Paraburkholderia atlantica* sp. nov. and *Paraburkholderia franconis* sp. nov., two new nitrogen-fixing nodulating species isolated from Atlantic forest soils in Brazil. Archives of Microbiology 202, 1369–1380. doi:10.1007/S00203-020-01843-W

Paulitsch, F., Dall’Agnol, R.F., Delamuta, J.R.M., Ribeiro, R.A., da Silva Batista, J.S., Hungria, M., 2019a. *Paraburkholderia guartelaensis* sp. nov., a nitrogen-fixing species isolated from nodules of *Mimosa gymnas* in an ecotone considered as a hotspot of biodiversity in Brazil. Archives of Microbiology 201, 1435–1446. doi:10.1007/S00203-019-01714-Z

Paulitsch, F., Delamuta, J.R.M., Ribeiro, R.A., da Silva Batista, J.S., Hungria, M., 2020b. Phylogeny of symbiotic genes reveals symbiovars within legume-nodulating *Paraburkholderia* species. Systematic and Applied Microbiology 43. doi:10.1016/J.SYAPM.2020.126151

Paulitsch, F., Klepa, M.S., da Silva, A.R., do Carmo, M.R.B., Dall’Agnol, R.F., Delamuta, J.R.M., Hungria, M., da Silva Batista, J.S., 2019b. Phylogenetic diversity of rhizobia nodulating native *Mimosa gymnas* grown in a South Brazilian ecotone. Molecular Biology Reports 46, 529–540. doi:10.1007/S11033-018-4506-Z

Peix, A., Ramírez-Bahena, M.H., Velázquez, E., Bedmar, E.J., 2015. Bacterial associations with legumes. Critical Reviews in Plant Sciences 34, 17–42. doi:10.1080/07352689.2014.897899

Platero, R., James, E.K., Rios, C., Iriarte, A., Sandes, L., Zabaleta, M., Battistoni, F., Fabiano, E., 2016. Novel *Cupriavidus* strains isolated from root nodules of native Uruguayan *Mimosa* species. Applied and Environmental Microbiology 82, 3150– 3164. doi:10.1128/AEM.04142-15

R Core Team, 2021. R: A language and environment for statistical computing. R Foundation for Statistical Computing. Vienna, Austria. URL https://www.R-project.org/.

Reeve, W., Ardley, J., Tian, R., Eshragi, L., Yoon, J.W., Ngamwisetkun, P., Seshadri, R., Ivanova, N.N., Kyrpides, N.C., 2015. A genomic encyclopedia of the root nodule bacteria: Assessing genetic diversity through a systematic biogeographic survey. Standards in Genomic Sciences 10. doi:10.1186/1944-3277-10-14

Remigi, P., Zhu, J., Young, J.P.W., Masson-Boivin, C., 2016. Symbiosis within symbiosis: evolving nitrogen-fixing legume symbionts. Trends in Microbiology 24, 63–75. doi:10.1016/J.TIM.2015.10.007

Richter, M., Rosselló-Móra, R., 2009. Shifting the genomic gold standard for the prokaryotic species definition. Proceedings of the National Academy of Sciences of the United States of America 106, 19126–19131. doi:10.1073/PNAS.0906412106

Sawana, A., Adeolu, M., Gupta, R.S., 2014. Molecular signatures and phylogenomic analysis of the genus *Burkholderia*: Proposal for division of this genus into the emended genus *Burkholderia* containing pathogenic organisms and a new genus *Paraburkholderia* gen. nov. harboring environmental species. Frontiers in Genetics 5. doi:10.3389/FGENE.2014.00429

Sheu, S.Y., Chou, J.H., Bontemps, C., Elliott, G.N., Gross, E., dos Reis Junior, F.B., Melkonian, R., Moulin, L., James, E.K., Sprent, J.I., Young, J.P.W., Chen, W.M., 2013. *Burkholderia diazotrophica* sp. nov., isolated from root nodules of *Mimosa* spp. International Journal of Systematic and Evolutionary Microbiology 63, 435–441. doi:10.1099/IJS.0.039859-0

Sheu, S.Y., Chou, J.H., Bontemps, C., Elliott, G.N., Gross, E., James, E.K., Sprent, J.I., Young, J.P.W., Chen, W.M., 2012. *Burkholderia symbiotica* sp. nov., isolated from root nodules of mimosa spp. native to north-east Brazil. International Journal of Systematic and Evolutionary Microbiology 62, 2272–2278. doi:10.1099/IJS.0.037408-0

Silva, F.C., 2009. Métodos de análises químicas para avaliação da fertilidade do solo. In: Silva F.C. (Ed). Manual de análises químicas de solos, plantas e fertilizantes: Embrapa Informação Tecnológica, 2009, pp. 107-184. http://www.infoteca.cnptia.embrapa.br/handle/doc/330496.

Silva, V.C., Alves, P.A.C., Rhem, M.F.K., dos Santos, J.M.F., James, E.K., Gross, E., 2018. Brazilian species of *Calliandra* Benth. (tribe Ingeae) are nodulated by diverse strains of Paraburkholderia. Systematic and Applied Microbiology 41, 241–250. doi:10.1016/J.SYAPM.2017.12.003

Simon, M.F., Grether, R., de Queiroz, L.P., Särkinen, T.E., Dutra, V.F., Hughes, C.E., 2011. The evolutionary history of *Mimosa* (Leguminosae): Toward a phylogeny of the sensitive plants1. American Journal of Botany 98, 1201–1221. doi:10.3732/AJB.1000520

Simon, M.F., Grether, R., De Queiroz, L.P., Skemae, C., Pennington, R.T., Hughes, C.E., 2009. Recent assembly of the Cerrado, a neotropical plant diversity hotspot, by in situ evolution of adaptations to fire. Proceedings of the National Academy of Sciences of the United States of America 106, 20359–20364. doi:10.1073/PNAS.0903410106

Simon, M.F., Proença, C., 2000. Phytogeographic patterns of *Mimosa* (Mimosoideae, Leguminosae) in the Cerrado biome of Brazil: An indicator genus of high-altitude centers of endemism? Biological Conservation 96, 279–296. doi:10.1016/S0006-3207(00)00085-9

Soares Neto, C.B., Ribeiro, P.R.A., Fernandes-Júnior, P.I., de Andrade, L.R.M., Zilli, J.E., Mendes, I.C., do Vale, H.M.M., James, E.K, dos Reis Junior, F.B., 2022. *Paraburkholderia atlantica* is the main rhizobial symbiont of *Mimosa* spp. in ultramafic soils in the Brazilian Cerrado biome. Plant and Soil 479, 465–479. doi:10.1007/S11104-022-05536-9

Sprent, J.I., 2009. Legume nodulation: A global perspective. Wiley-Blackwell, USA. doi:10.1002/9781444316384

Sprent, J.I., Ardley, J., James, E.K., 2017. Biogeography of nodulated legumes and their nitrogen-fixing symbionts. New Phytologist 215, 40–56. doi:10.1111/NPH.14474

Stopnisek, N., Bodenhausen, N., Frey, B., Fierer, N., Eberl, L., Weisskopf, L., 2014. Genus-wide acid tolerance accounts for the biogeographical distribution of soil *Burkholderia* populations. Environmental Microbiology 16, 1503–1512. doi:10.1111/1462-2920.12211

Taulé, C., Zabaleta, M., Mareque, C., Platero, R., Sanjurjo, L., Sicardi, M., Frioni, L., Battistoni, F., Fabiano, E., 2012. New betaproteobacterial rhizobium strains able to efficiently nodulate *Parapiptadenia rigida* (Benth.) Brenan. Applied and Environmental Microbiology 78, 1692–1700. doi:10.1128/AEM.06215-11

Trifinopoulos, J., Nguyen, L.-T., von Haeseler, A., Minh, B.Q., 2016. W-IQ-TREE: a fast online phylogenetic tool for maximum likelihood analysis. Nucleic Acids Research 44, W232–W235. doi:10.1093/nar/gkw256

Vincent, J.M., 1970. A manual for the practical study of root-nodule bacteria. Oxford: Blackwell Scientific (International Biological Programme handbook, 15), 1970.

Vinuesa, P., Silva, C., Lorite, M.J., Izaguirre-Mayoral, M.L., Bedmar, E.J., Martínez-Romero, E., 2005. Molecular systematics of rhizobia based on maximum likelihood and Bayesian phylogenies inferred from *rrs*, *atpD*, *recA* and *nifH* sequences, and their use in the classification of *Sesbania* microsymbionts from Venezuelan wetlands. Systematic and Applied Microbiology 28, 702–716. doi:10.1016/j.syapm.2005.05.007

Zilli, J.É., de Moraes Carvalho, C.P., de Matos Macedo, A.V., de Barros Soares, L.H., Gross, E., James, E.K., Simon, M.F., de Faria, S.M., 2021. Nodulation of the neotropical genus *Calliandra* by alpha or betaproteobacterial symbionts depends on the biogeographical origins of the host species. Brazilian Journal of Microbiology 52, 2153–2168. doi:10.1007/s42770-021-00570-8

